# Drought influences the fungal community structure, diversity, and functionality inhabiting the grapevine xylem and enhances the abundance of *Phaeomoniella chlamydospora*

**DOI:** 10.1101/2024.02.28.582583

**Authors:** C. Leal, R. Bujanda, M. J. Carbone, T. Kiss, A. Eichmeier, D. Gramaje, M. M. Maldonado-González

## Abstract

The productivity of grapevines in Mediterranean regions faces significant threats from global warming, which may intensify competition for water resources. Recent research highlights the impact of water deficit on the root-associated microbiota of grapevines, particularly organisms capable of mitigating abiotic and biotic stressors. This study explores the influence of drought on the structure, diversity, and functionality of xylem-inhabiting fungal communities of grapevine, with a focus on the fungal pathogen *Phaeomoniella chlamydospora* associated with esca and Petri diseases. One-year-old grapevine rootlings grown under greenhouse conditions were subjected to three water regimes: severe water deficit (SWD) at 25% of field capacity, moderate water deficit (MWD) at 50% of field capacity, and no water deficit (AWD) at 100% of field capacity. Wood samples were non-destructively collected before planting (t0) and one (t1) and two (t2) growing seasons after planting from the bottom, medium, and apical parts of the rootstock. Fungal composition and *P. chlamydospora* abundance were assessed using ITS high-throughput amplicon sequencing (HTAS) and droplet-digital PCR (ddPCR), respectively. The induced water stress not only altered the diversity and composition of the fungal microbiome in the xylem vessels but also affected co-occurrence networks, resulting in less complex networks with fewer correlations between taxa, potentially increasing grapevine vulnerability to various biotic and abiotic stresses. SWD significantly reduced microbial diversity, leading to a shift in the abundance of pathotrophs such as *P. chlamydospora* in the xylem. This underscores the interconnectedness between water stress, microbiome dynamics, and plant health. The combination of compromised plant defenses, altered physiological conditions, and shifts in the surrounding microbial community may create conditions conducive to increased *P. chlamydospora* abundance in the xylem vessels of young vines following water stress.

## INTRODUCTION

Climate change is one of the most defining concerns of today’s world and has greatly reshaped or is in the process of altering all agricultural ecosystems (Arora, 2019). In viticulture, the major consequence of climate change is the higher risk of water stress (drought) (Jones et al. 2005; Schultz and Stoll, 2010; Mosedale et al. 2016; Van Leeuwen and Darriet, 2016). Grapevine growing regions worldwide have recently faced intense and frequent droughts and heat waves, e.g. 2009 in Australia, 2015 in California, and 2019 in France, and subsequent economical losses in wine production have been considerable (Lamarque et al. 2023). Grapevine is mostly grown in Mediterranean climate regions where seasonal drought and temperature exert large constraints on yield and quality (Chaves et al. 2010, Mosedale et al. 2016).

Drought exerts significant impacts on grapevines, influencing their growth, physiology, and overall health (Gambetta et al., 2020), thereby shaping the development and progression of grapevine diseases. In certain instances, drought stress can exacerbate disease severity and hasten disease progression, while in others, it may suppress pathogen growth and mitigate disease severity. Moreover, the specific consequences of water scarcity appear to be contingent not only on the pathogen but also on prevailing environmental conditions (Liu et al., 2019). In terms of defense mechanisms, stressed plants often experience a reduction in these mechanisms, rendering them more susceptible to pathogen infections (Pérez-Harguindeguy et al., 2016).

Among the most significant fungal diseases affecting grapevines are grapevine trunk diseases (GTDs), caused by various pathogens, which can be influenced by drought stress (Sosnowski et al., 2016). These diseases pose a substantial threat to vineyard sustainability and exert a detrimental impact on plant productivity, serving as a primary contributor to vine decline, with potential long-term consequences such as plant mortality (Gramaje et al., 2018). Drought stress emerges as arguably the most critical factor in the manifestation of GTD symptoms, thus it comes as no surprise that it has been extensively investigated among all potential stress factors (Fischer and Kassemeyer, 2012; Scala et al., 2019; Bortolami et al., 2019, 2021).

Esca is a complex GTD known to occur wherever grapevines are grown, whose symptoms have been shown to result not only from a biological agent but also from physiological factors affecting the framework of the vine (Bortolami et al. 2023). Esca is associated with mature grapevines and presents in two forms: the slow form, characterized by foliar symptoms known as ’tiger-stripe,’ and the fast form, characterized by sudden plant death (Lecomte et al. 2012). Petri disease is associated with the young vine decline syndrome, resulting in lower yield, poor vigor, and premature death of grapevines, usually in field nurseries and within five years after planting (Gramaje and Armengol, 2011). In both diseases, internal symptoms are characterized by necrotic spots in the xylem, primarily isolating the ascomycete species *Phaeomoniella chlamydospora* (Gramaje et al. 2018).

To bolster resistance against microbial pathogens and enhance tolerance to abiotic stressors like drought, grapevines form symbiotic relationships with diverse communities of microorganisms that also facilitate plant growth (Hacquard et al., 2017; Hardoim et al., 2015; McLaren and Callahan, 2020; Ngumbi and Kloepper, 2016; Pineda et al., 2017). These plant-associated microbiomes showcase remarkable diversity and complexity, yet our comprehension of the mechanisms and factors governing specific plant-associated microbial communities remains limited (Reinhold-Hurek et al., 2015).

The objective of this study has been to investigate alterations in the xylem-inhabiting fungal communities of grapevine following the application of water deficit, with particular attention to the primary fungal pathogen associated with esca and Petri diseases, *P. chlamydospora* (Gramaje et al. 2018). We focused on drought, one of the abiotic stresses that is postulated to increase in occurrence and severity across the globe and have direct significant impacts on grapevine productivity and ecosystem sustainability (Gambetta et al. 2020).

## MATERIALS AND METHODS

### Experimental design and treatments

In April 2017, 144 dormant grapevine plants of Tempranillo cultivar grafted onto rootstock 110 Richter (110 R) were obtained from a commercial nursery located in Lagarra (northern Spain). The grapevines were individually planted in 11-liter pots, each pot housing one plant. The pots were filled with peat substrate, with a 2-centimeter layer of vermiculite positioned at the bottom. Plants were grown in a greenhouse at the Institute of Grapevine and Wine Sciences (ICVV) (42C:26’39.512” N 2C:30’51.229” W) from 27 April 2017 to 1 December 2018.

One month after sprouting, potted plants were randomly divided into three treatments, simulating three types of irrigation regimes. The treatments were (1) irrigation at 25% of field capacity (severe water deficit: SWD), (2) irrigation at 50% of field capacity (moderate water deficit: MWD) and (3) irrigation at 100% of field capacity (no water deficit: AWD). Soil water content at field capacity was previously calculated according to O’Geen (2013). An automated drip irrigation system was adjusted for each irrigation treatment, by measuring the dielectric constant of soil. The irrigation treatments were maintained from May 2017 to November 2017, and from March 2018 to November 2018. During the winter (December 2017-February 2018), the irrigation was maintained constant in all scenarios. The experimental design consisted of two randomized blocks per irrigation treatment, each containing 12 plants (24 plants per irrigation treatment). The experiment was repeated in two greenhouse rooms (72 plants per room). In both greenhouse rooms, plants were irrigated with nutritive solution (0.1 mM NH_4_H_2_PO_4_, 0.187 mM NH_4_NO_3_, 0.255 mM KNO_3_, 0.025 mM MgSO_4_, 0.002 mM Fe, and oligoelements [B, Zn, Mn, Cu, and Mo]); climatic conditions were monitored every 15 min using temperature and humidity probes (S-THB-M002, Onset) and global radiation sensors (S-LIx-M003, Onset) connected to a data logger (U300- NRC, Onset).

Stomatal conductance was evaluated using a Decagon SC-1 porometer (AlphaOmega Electronics, Spain) with an approximate frequency of 15 days at solar noon (Oliveira et al. 2014). For this purpose, measurements were taken from 11 plants for each water regime in July 2017 for the first growing season, and from May to August 2018 during the second growing season. Predawn leaf water potential (LWP) was measured with a pressure chamber (PMS Instrument, USA) (Scholander et al. 1965). During the first growing season, the measurement of LWP was conducted once on 5 plants per water regime on October 26, 2017. During the second growing season, monitoring of water potential was carried out every 15 days from May 2018 to August 2018. At the end of the second growing season, plants were assessed for undried shoot weight.

The data were subjected to analysis of variance and mean values were separated according to Tukeýs honestly significant difference at *P* = 0.05, with Statistix 10 software (Analytical Software, FL, USA).

### Sample collection

A 0.5 mm micro drill, specifically the MICROMOT 50/EF (Proxxon micromot, Madrid, Spain), was utilized for the extraction of grapevine wood samples from the xylem vessels of 110 R rootstock (Gramaje et al. 2022). Woody tissues were procured from three distinct sections of each rootstock cutting: the base (1 cm above the basal part of the cutting), the middle, and the apical (1 cm below the graft union). This sampling was conducted at three specific time points: prior to planting in April 2017 (t0; 10 plants), nine months later in December 2017 (t1; 12 plants for each water regime and room), and 21 months later in December 2018 (t2; the remaining 12 plants for each water regime and room) subsequent to the initiation of the experiment. For the sampling process, the plants were uprooted and longitudinally bisected using sterile scissors. From each section of the plant, 20 mg of woody tissue were harvested and placed into sterile Eppendorf tubes, accumulating a total of 60 milligrams mg of tissue per plant.

### DNA extraction, sequencing and data analysis of the high-throughput amplification assay

DNA was obtained from the xylem tissue collected during each sampling period using the i-genomic Plant DNA Extraction Mini Kit (Intron Biotechnology, Seongnam, KR). To measure the DNA yields from each sample, we utilized the Invitrogen Qubit 4 Fluorometer with the Qubit dsDNA HS Assay (Thermo Fisher Scientific, Waltham, USA), and the resulting extracts were adjusted to a concentration of 10 ng/μl. Samples collected from the three different sections of each rootstock cutting were pooled resulting in a single sample per plant. A total of 154 DNA samples was analyzed (10 samples at t0, 72 samples at t1, and 72 samples at t2). The primers ITS86F (5’ GTGAATCATCGAATCTTTGAA 3’) (Turene et al. 1999) and ITS4 (5’TCCTCCGCTTATTGATATGC 3’) (White et al. 1990) were used to amplify the complete fungal ITS2 region (around 300 bp). Illumina sequencing primer sequences were attached to their 5’ ends.

PCRs were conducted in a final volume of 25 µL, comprising 12.5 µL of Supreme NZYTaq 2x Green Master Mix (NZYTech, Lisboa, Portugal), 0.5 µM of the primers, 2.5 µL of template DNA, and ultrapure water to reach a total volume of 25 µL. The following PCR protocol was applied: an initial denaturation step at 95 °C for 5 minutes, followed by 35 cycles of 95 °C for 30 seconds, 49 °C for 30 seconds, 72 °C for 30 seconds, and a final extension step at 72 °C for 10 minutes. During a second round of PCR, the oligonucleotide indices were attached using the same conditions. However, for an overview of the library preparation process, only five cycles were carried out, and the annealing temperature was set at 60 °C. To monitor contamination during library preparation in each PCR round, a negative control with no DNA was included. Additionally, a positive control consisted of DNA from a grapevine wood sample previously assessed by ITS2 High throughput amplicon sequencing (HTAS) (Martínez- Diz et al. 2020). Library sizes were confirmed using 2% agarose gels stained with GreenSafe (NZYTech, Lisboa, Portugal). The libraries were subsequently purified using Mag-Bind RXNPure Plus magnetic beads (Omega Biotek, Norcross, GA, USA), and then combined in equimolar amounts based on the quantification data obtained from the Qubit dsDNA HS Assay (Thermo Fisher Scientific, Waltham, USA). The resulting pool was subjected to sequencing in a MiSeq PE300 run (Illumina, San Diego, USA). Control samples were sequenced to assess the potential for contamination during the process.

The quality assessment of the sequencing data was carried out using FastQC-0.10.1. Subsequent data processing was performed using SEED v2.1.2 (Vetrovsky et al. 2018). The raw forward and reverse sequences for each sample were merged into paired-end reads using the fastq-join 1.1.2 tool from the ea-utils suite (Aronesty 2011). After that, the sequences underwent quality filtering, with a threshold of Q = 30, and removal of sequences with less than 250 bases. Sequences with any ambiguous bases were removed as well. The sequences were then grouped based on their corresponding sample names. Fungal ITS sequences were extracted using ITSx 1.0.11 (Bengtsson-Palme et al. 2013). Following this, the sequences were clustered into operational taxonomic units (OTUs) using UPARSE implementation in USEARCH ver. 8.1.1861 (Edgar 2013) with 97% similarity threshold. In this step, the chimeric sequences were identified and removed. The representative most abundant sequence of each OTU was identified using MAFFT 7.222 (Katoh et al. 2009). Finally, the identification of OTUs was performed using blastn, tblastx, and makeblastdb 2.5.0+ (https://blast.ncbi.nlm.nih.gov/Blast.cgi, accessed on 28 May 2023) against local UNITE 8.3 fungal dynamic database (Abarenkov et al. 2010). To ensure comparability between samples, the dataset was normalized using the Total Sum Scaling standard approach.

The OTU table, metadata and taxonomic classifications used in this study have been deposited in Figshare (ID: 197131). HTAS data have been deposited in GenBank/NCBI under BioProject Acc. No. PRJNA1078128.

### Fungal diversity, taxonomy distribution and statistical analysis

Alpha diversity metrics were calculated utilizing the Shannon and Chao1 indices within the Phyloseq package, facilitated by the MicrobiomeAnalyst 2.0 software (Lu et al. 2023). To ensure adequate data coverage and construct rarefaction curves, the MicrobiomeAnalyst tool was employed. Discriminant analysis of taxa relative abundance (at the genus level or higher) across different water stress regimes and sampling times was performed using the Linear Discriminant Analysis Effect Size (LEfSe) algorithm (Segata et al. 2011), implemented through MicrobiomeAnalyst. A logarithmic Linear Discriminant Analysis (LDA) score threshold of 2.0 and a False Discovery Rate (FDR)-adjusted *P* value cutoff of 0.1 were set. Shared fungal Operational Taxonomic Units (OTUs) across water stress regimes and sampling times were identified through Venn diagram analysis using software accessible at http://bioinformatics.psb.ugent.be.

Correlation network analysis at the genus taxonomic level was executed using MicrobiomeAnalyst, employing the SparCC algorithm (Friedman and Alm, 2012) with 100 permutations, a *P* value threshold of 0.01, and a correlation threshold of 0.5. Co- occurrence network analysis, aimed at exploring potential interactions between genera, was conducted using the integrated Network Analysis Pipeline (iNAP) (Feng et al. 2022), setting a *P* value threshold of 0.05 with 120 permutations, and a correlation threshold of 0.3 to identify significant associations. The resultant network was visualized using Cytoscape version 3.10.0 (Shannon et al. 2003).

### Functional prediction of fungal communities

We investigated the role of fungal communities within the xylem vessels under three different irrigation conditions using FUNGuild v1.0, as described by Nguyen et al. (2016). Eleven guilds were classified based on three trophic modes: pathotrophs, saprotrophs, and symbiotrophs. These guilds included plant pathogens, animal pathogens, fungal parasites, lichen parasites, undefined saprotrophs, soil saprotrophs, wood saprotrophs, dung saprotrophs, plant saprotrophs, endophytes, and arbuscular mycorrhizal fungi. OTUs that did not match taxa in the database were categorized as "unassigned." Guilds classified as "probable" and "highly probable" according to the fungal database were selected for further analysis.

We calculated the relative abundance of OTUs based on these guilds in the xylem vessels under the three irrigation conditions. To assess the impact of water stress conditions on the relative abundance of OTUs within different trophic modes, we conducted ANOVA using Statistix 10 software (Analytical Software). Prior to analysis, the data were transformed to the square root (√x). We compared the means of transformed data using Tukey’s honestly significant difference test at a significance level of *P* = 0.05.

### Droplet digital PCR assay

To quantify the natural inoculum of *P. chlamydospora* in rootstock cuttings, we conducted droplet digital PCR (ddPCR) analyses using DNA extracted from the rootstock wood. This analysis utilized specific primers and a probe assay designed to target the beta-tubulin region (Hrycan et al. 2023). Each reaction consisted of 750 nM of each primer, 1x Supermix for Probes (Bio-Rad, Hercules, CA, USA), 250 nM of the probe, and 2 µl of DNA, making a total volume of 20 µl. Droplets were generated using the Bio-Rad QX200^TM^ droplet generator (Bio-Rad, Hercules, CA, USA), with the entire reaction volume per sample and 70 µl of QX200^TM^ droplet generation oil.

The PCR runs were performed in a Bio-Rad C1000 touch thermal cycler (Bio-Rad, Hercules, CA, USA) with the following cycling conditions: an initial heating step at 95°C for 10 minutes, followed by 40 cycles of denaturation at 94°C for 30 seconds, annealing at 55°C for 60 seconds, and a final incubation at 98°C for 10 minutes. The PCR plates were then transferred to the Bio-Rad QX200^TM^ droplet reader (Bio-Rad, Hercules, CA, USA), and the results were analyzed using QuantaSoftTM software (Bio- Rad, Hercules, CA, USA). For the analysis, DNA from the *P. chlamydospora* isolate BV-0577 was used as a template. The threshold was manually set at 3,000, based on two positive controls, one containing DNA from a pure culture of *P. chlamydospora* isolate BV-0577 and the other containing DNA from a grapevine wood sample previously identified as positive for *P. chlamydospora* using ITS HTAS. Additionally, a non-template control (NTC) reaction (using water) was included in the analysis. All samples were run in triplicate by ddPCR. Following Bio-Rad’s recommendation (https://www.bio-rad.com/webroot/web/pdf/lsr/literature/Bulletin_6407.pdf), wells with fewer than three positive droplets were considered negative.

### Statistical analysis

The *P. chlamydospora* read counts were subjected to a logarithmic transformation using the formula log (n/N * 1000 + 1), where ’n’ represented the number of reads detected in each sample, and ’N’ was the total number of reads detected. Additionally, DNA concentration, expressed as copies per µl, was log-transformed before analysis. To assess differences in HTAS abundance and ddPCR copy numbers among the various sampling moments, a one-way ANOVA test was employed with the assistance of Statistix 10 software (Analytical Software, FL, USA). Subsequently, Tukey’s honestly significant difference test was applied to compare the means of the data (*P* = 0.05). Furthermore, correlation analysis between the transformed datasets from both HTAS and ddPCR was conducted using the corrr package in R version 4.4 (R Core Team, 2023).

## RESULTS

### Stomatal conductance, water potential and shoot weight

Stomatal conductance was overall maintained during time from the beginning of the experiment to the end of year 2 (Fig. S1). Plants with AWD maintained a stomatal conductance between 272 and 243 mmol m^-2^ s^-1^ during the first growing season, and between 333 and 270 mmol m^-2^ s^-1^ during the second growing season. Plants with MWD had values that range from 204 and 148 mmol m^-2^ s^-1^ during the first growing season, and between 212 and 176 mmol m^-2^ s^-1^ during the second growing season, and plants with SWD presented the lowest stomatal conductance values ranging from 121 and 101 mmol m^-2^ s^-1^ during the first growing season, and between 133 and 108 mmol m^-2^ s^-1^ during the second growing season.

During the first growing season, the LWP of plants with AWD varied from -0.6 to -0.7 Mpa, plants with MWD varied from -0.8 to -0.9 Mpa, and plants with SWD varied from -0.75 to -0.9 Mpa (Fig. S1). During the second growing season, the LWP of plants with AWD varied from -0.45 to -0.61 Mpa, plants with MWD varied from -0.76 to -0.91 Mpa, and plants with SWD varied from -1.27 to -1.59 Mpa. The variable irrigation regime had a significant effect on the shoot weight (*P* < 0.01) (Fig. S2). Treatment AWS (176.0 g ± 5.7) differed significantly from the MWD (91.9 g ± 2.7) and SWD (57.1 g ± 3.1) treatments.

### Sequencing depth and community diversity

After paired-end alignments, quality filtering and deletion of chimeras, singletons, a total of 1,853,314 fungal ITS2 sequences were generated from 150 samples (four samples were removed from the analysis due to the low number of reads), and assigned to 238 fungal operational taxonomic units (OTUs). According to the Good’s coverage values, 99.94% of the total species richness were accounted for in fungal communities. All diversity was captured with an adequate sequencing depth (Fig. S3). Chao1 richness estimator ranged from 21.61 to 24.11 during the first growing season, and from 17.02 to 20.02 during the second growing season. Shannon diversity estimator ranged from 2.19 to 2.31 during the first growing season, and from 1.91 to 1.97 during the second growing season (Table S1).

### Water deficit affects fungal diversity in both sampling times

There were no significant differences in alpha diversity between plant stocks obtained from the two greenhouse rooms for each of the sampling times (*P* > 0.05); therefore, the data from the two plant stocks were analyzed together for each sampling time. Our results showed significant diversity differences between sampling times (Chao1 index <0.01; Shannon index: 0.0306) (Fig. S4), therefore, all analyses were performed for each sampling time separately. Fungal microbiome varied significantly among irrigation regimes (Fig. 1), with the exception of the Shannon index in t2. Samples with AWD presented the highest alpha-diversity index (t1: Chao1 24.11 ± 1.07 and Shannon 2.31 ± 0.08; t2: Chao1 20.02 ± 0.45 and Shannon 1.97 ± 0.03), followed by the MWD treatment (t1: Chao1 21.92 ± 1.98 and Shannon 2.19 ± 0.10; t2: Chao1 18.57 ± 0.48 and Shannon 1.99 ± 0.04). The treatment with the lowest alpha-diversity was SWD (t1: Chao1 21.61 ± 1.07 and Shannon 2.22 ± 0.03; t2: Chao1 17.02 ± 1.41 and Shannon 1.91 ± 0.06), where plants present higher water stress (Table S1).

**Fig. 1.**
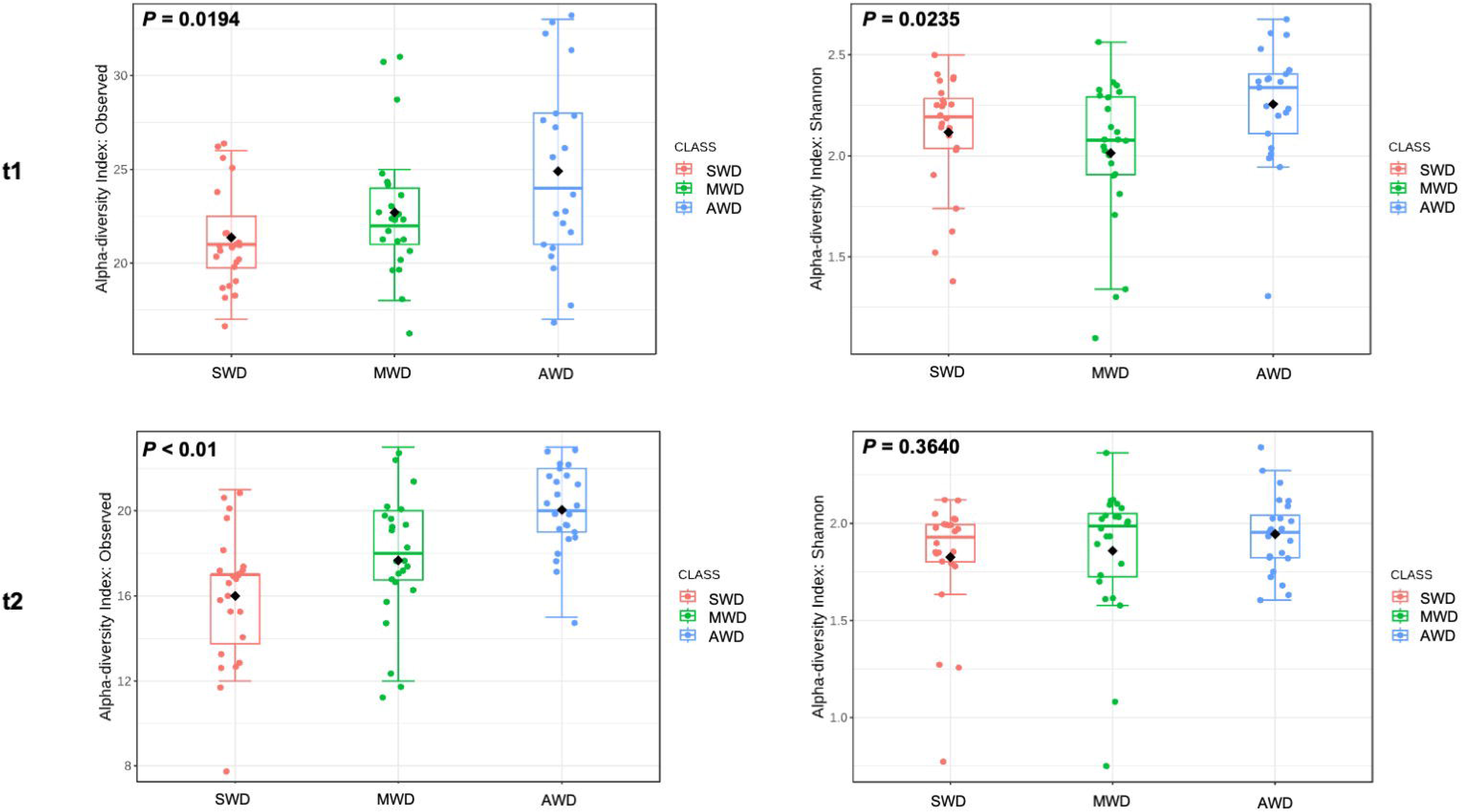
Alpha diversity of fungi in the xylem microbiome following one (t1) and two (t2) growing seasons: comparative analysis among Severe Water Deficit (SWD), Moderate Water Deficit (MWD), and No Water Deficit (AWD) treatments. P-values are indicated within the graphs.

### Irrigation Regime-Specific and Shared Fungal Assemblages

For each sampling time, the three water regimes showed specific fungal OTUs and a cluster of shared OTUs (Fig. 2). In t1, 40.3% (77 OTUs) of fungal OTUs were shared among irrigation regimes. Specific fungal OTUs differed according to irrigation regime, with 13.6% (26 OTUs) for SWD, 9.4% (18 OTUs) for MWD, and 12.6% (24 OTUs) for AWD. In t2, 37.0% (71 OTUs) of fungal OTUs were shared among irrigation regimes. Specific fungal OTUs differed according to irrigation regime, with 9.9% (19 OTUs) for SWD, 13.0% (25 OTUs) for MWD, and 20.8% (40 OTUs) for AWD.

**Fig. 2.**
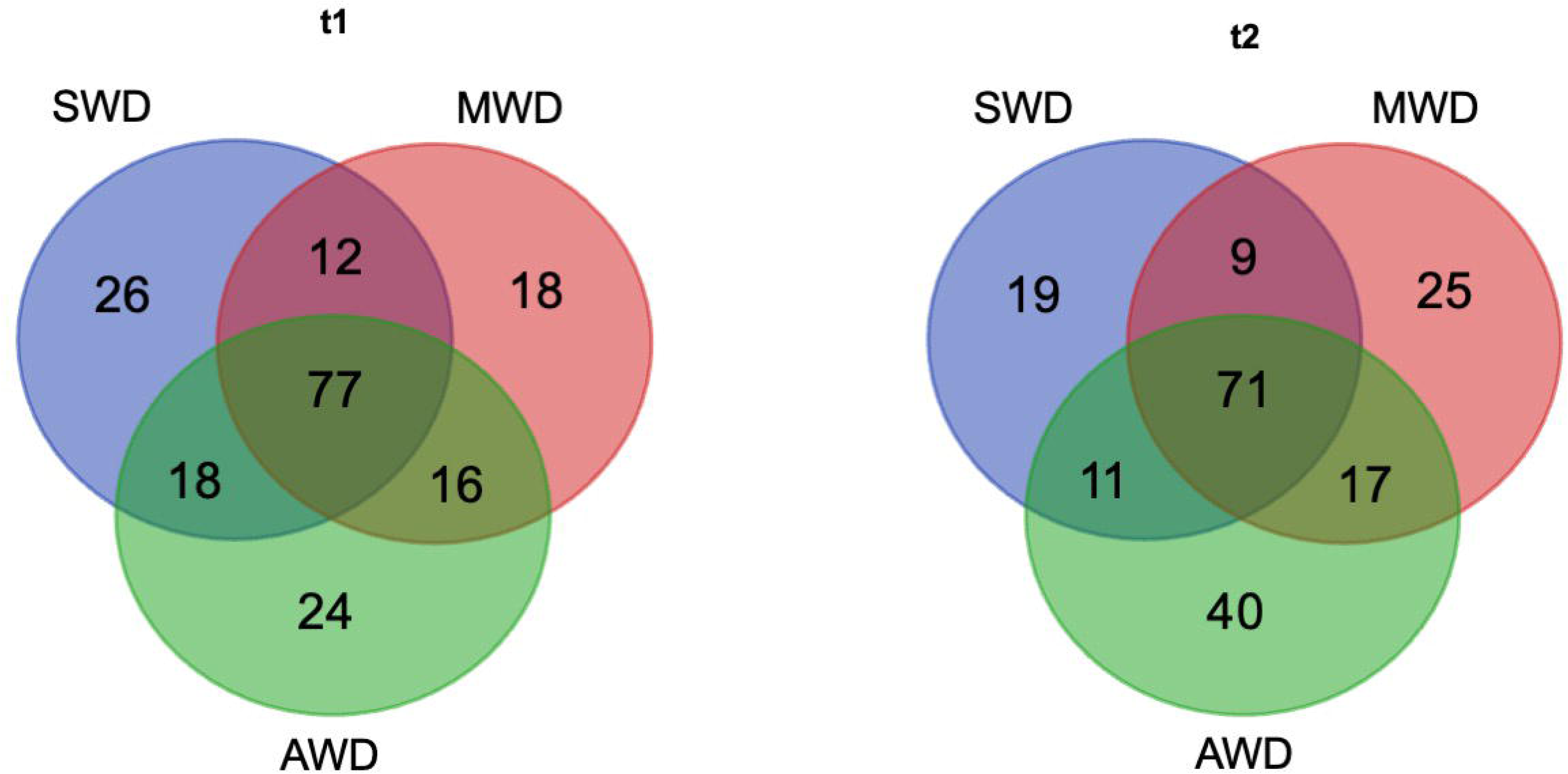
Venn diagram depicting the shared and unique Operational Taxonomic Units (OTUs) in the fungal microbiota after one (t1) and two (t2) growing seasons across treatments: Severe Water Deficit (SWD), Moderate Water Deficit (MWD), and No Water Deficit (AWD).

The composition of the microbiome was generally maintained over time (Fig. 3). The microbiome consisted majorly in *Acremonium, Monocillium*, and *Penicillium*, although at t2, the major genus was *Wallemia*. Other genera like *Minimelanolocus, Calyptosphaeria, Phlogicylindrium, Talaromyces, Ceratobasidium, Malassezia*, and *Sporothrix* were also found in the xylem microbiome, but in less quantity than the others.

**Fig. 3.**
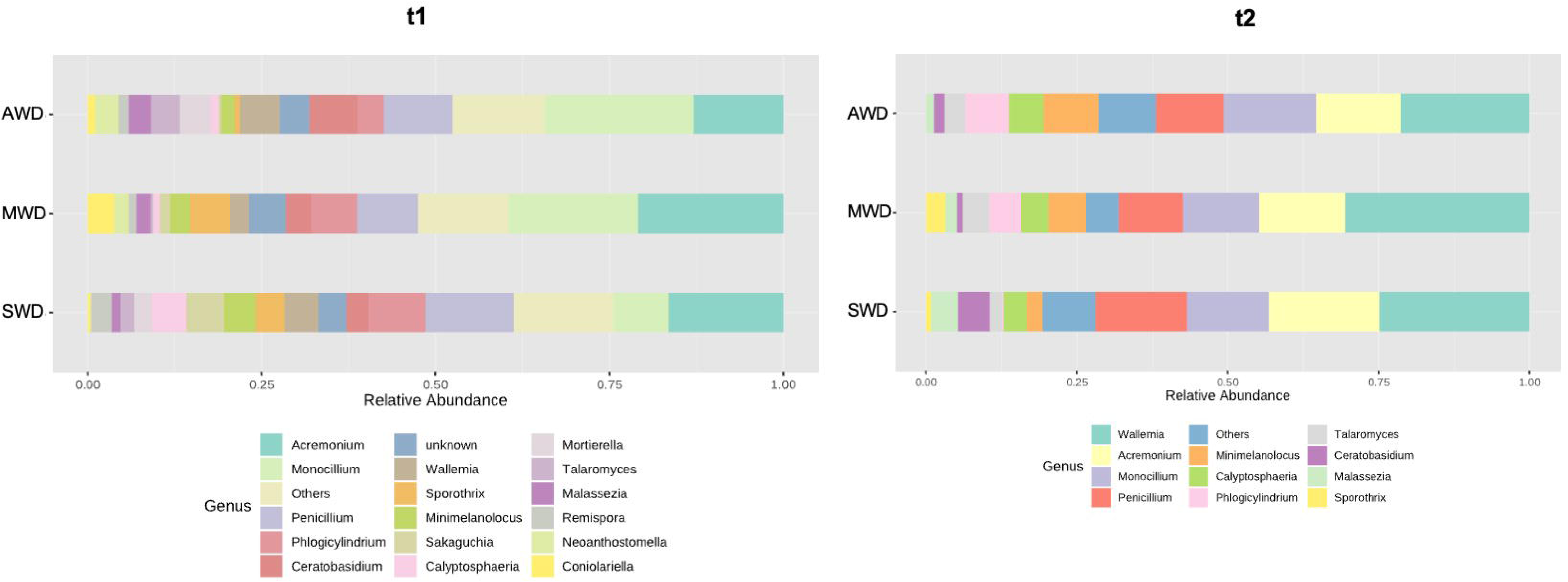
Fungal microorganism abundance in the xylem microbiome following one (t1) and two (t2) growing seasons. Treatments include Severe Water Deficit (SWD), Moderate Water Deficit (MWD), and No Water Deficit (AWD).

Linear discriminant analysis Effect Size (LEfSe) analysis determined 10 genera most likely to explain significant differences between irrigation regimes at t1, and 15 at t2 (Fig. 4). At t1, all genera were more abundant in plants with AWD, except for *Phaeomoniella* and *Wallemia* that were more abundant in plants with SWD, and *Cadophora* that was the most abundant in plants with MWD. At t2, all genera were more abundant in plants with AWD, except for *Acremonium*, *Ceratobasidium*, *Malassezia*, and *Phaeomoniella* that were more abundant in plants with SWD, and *Wallemia* and an unkown group that were most abundant in plants with MWD.

**Fig. 4.**
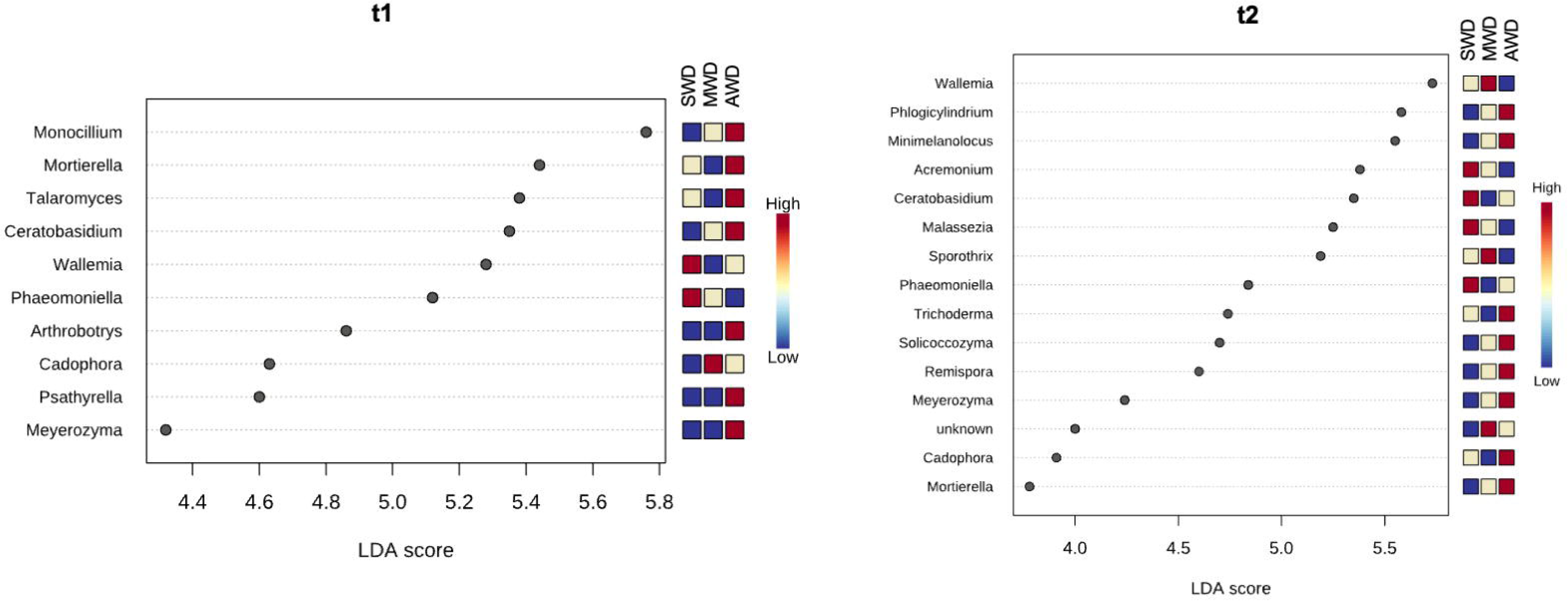
Linear Discriminant Analysis Effect Size (LEfSe) for different treatments following one (t1) and two (t2} growing seasons: Severe Water Deficit (SWD), Moderate Water Deficit (MWD), and No Water Deficit (AWD).

### *Phaeomoniella* correlated negatively with pathogens and biocontrol agents

A similar number of significant edges and connections was observed at both sampling times (t1: n = 82, t2: n = 84) (Fig. 5; Table S2). At t1, *Phaeomoniella* correlated positively with *Phlogicylindrium* and negatively with *Cadophora* and *Psathyrella*. Other pathogens such as *Cadophora* correlated positively with *Monocillium* and negatively with *Penicillium* and *Sakaguchia*, while *Phaeoacremonium* correlated positively with *Arthrobotrytis* and *Lyomeces*. At t2, *Phaeomoniella* correlated positively with *Ceratobasidium* and *Wallemia*, and negatively with *Phlogicylindrium* and the biological control agent *Trichoderma*. Furthermore, *Trichoderma* also showed a negative correlation with *Sakaguchia*. *Cadophora* correlated positively with *Solicoccozyma* and negatively with *Sporothrix*, while *Phaeoacremonium* correlated positively with *Talaromyces* and negatively with *Solicoccozyma*.

**Fig. 5.**
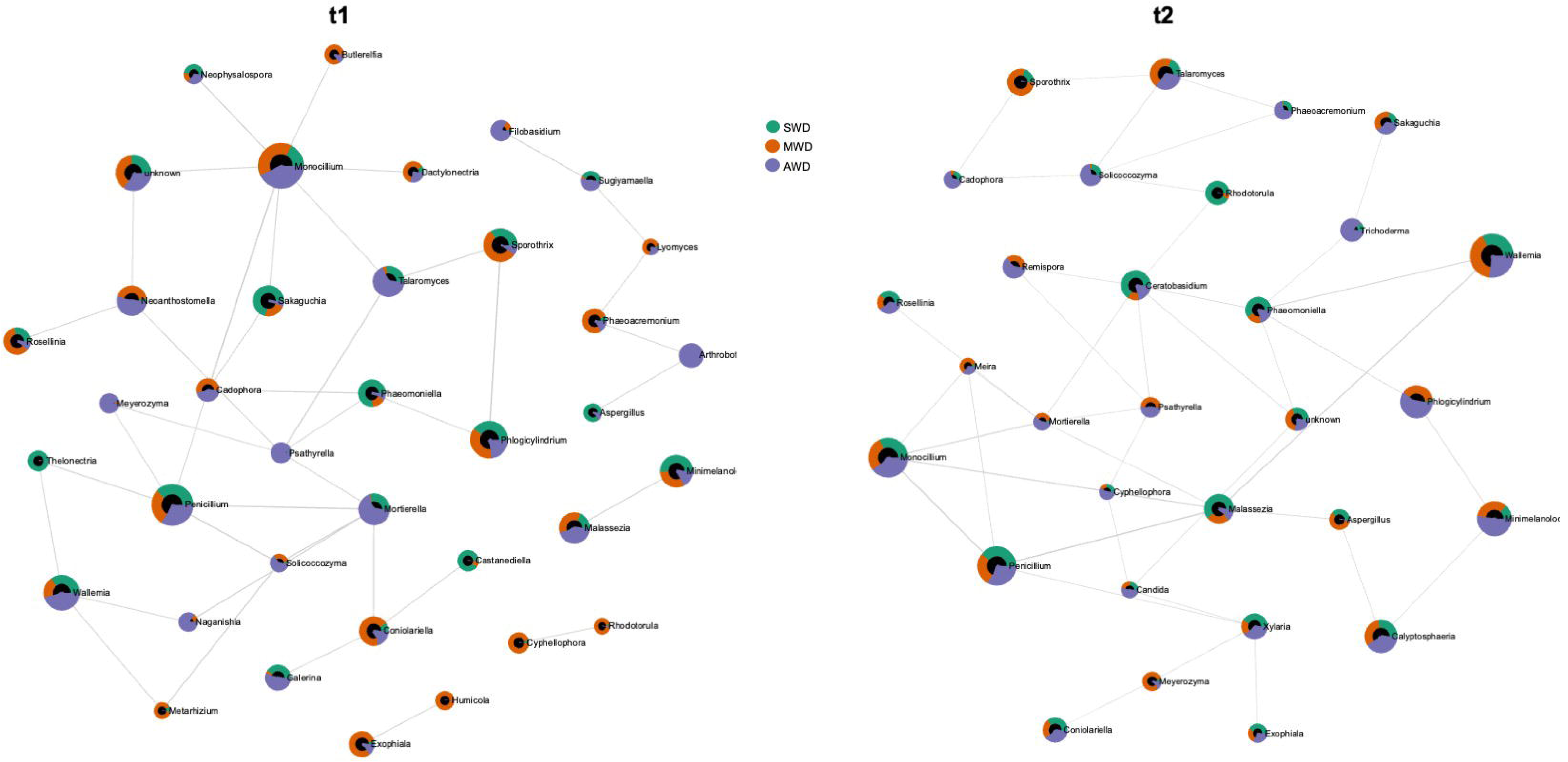
SparCC correlation analysis at genus level across irrigation regimes (SWD: severe water deficit, MWD: moderate water deficit and AWD: no water deficit) after one (t1) and two (t2) growing seasons

### Water deficit affects fungal functionality

Overall, the relative abundance of fungal OTUs identified as trophic modes with pathotrophs, saprotrophs, and symbiotrophs ranged from 93.3% to 94.4% at t1, and 91.0% to 92.7% at t2, while the remaining OTUs were unassigned (Fig. 6). There were significant differences in the relative proportion of fungal functions within each irrigation regimes in each sampling time (*P* < 0.05) (Table S3).

**Fig. 6.**
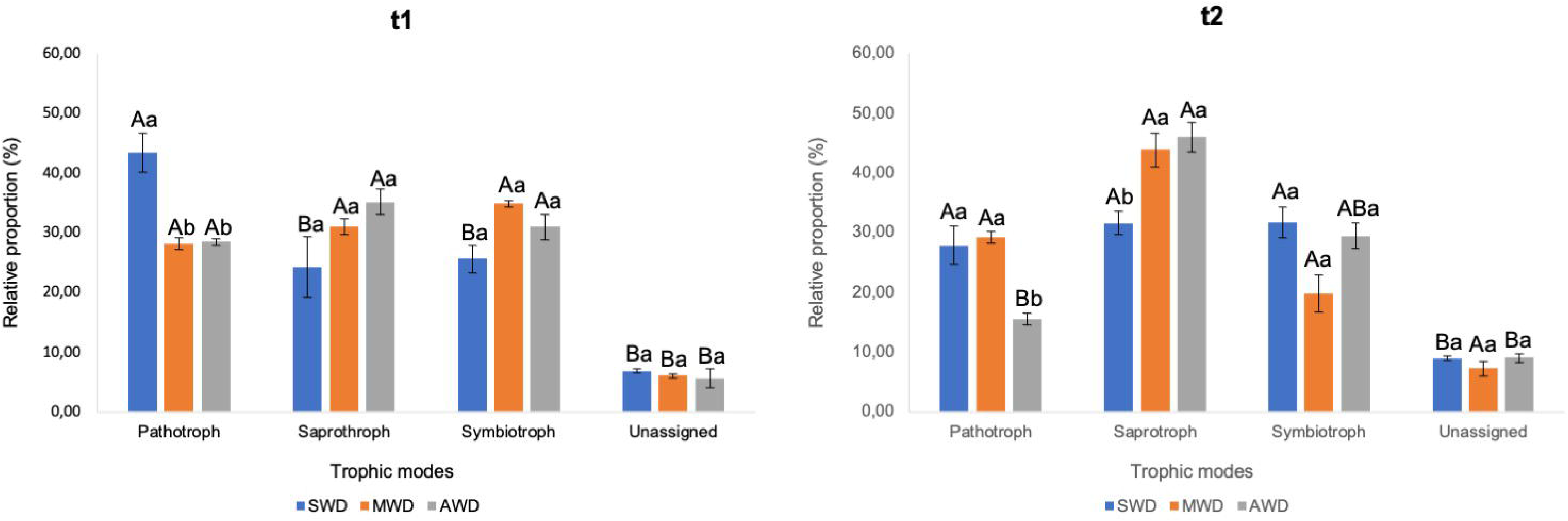
Variation in fungal functions analyzed by FUNGuild across growing seasons and irrigation regimes: a comparative study over two Seasons (t1 and t2) under Severe Water Deficit (SWD), Moderate Water Deficit (MWD), and No Water Deficit (AWD) conditions. Statistical significance assessed with Tukey’s Test at p < 0.05. Capital letters are for comparison of means among functional groups within each irrigation regime. Small letters are for comparison of means among irrigation regimes within each functional group.

At t1, the relative proportion of pathotrophs was significantly higher in SWD (43.5%) than in MWD (28.1%) and AWD (28.4%) (Fig. 6; Table S3). At t2, the relative proportion of pathotrophs was significantly higher in SWD (27.8%) and MWD (29.1%) than AWD (15.5%). Saprotrophs were found at SWD treatment (31.5%) in lower proportion compared with MWD (43.9%) and AWD (45.9%).

At both sampling times, plant pathogens were the dominant taxa in the pathotroph group in all water regimes (Table S4). At t1, there were significant differences in plant pathogen abundances between SWD (31.2%) and AWD (12.3%), while at t2 the significant differences were found between MWD (18.9%) and AWD (8.6%). In the saprotroph group, undefined saptrotrophs were the dominant taxa in the three water regimes at both sampling times, except for AWD at t2 in which wood saprotrophs were the dominant taxa (Table S4). The symbiotroph group was dominated by endophytes in all water regimes at both sampling times.

### Water deficit increases the absolute abundance and sequencing reads of *P. chlamydospora*

The analysis showed a positive significant correlation between the number of OTUs and the *P. chlamydospora* DNA quantified using ddPCR (*P* < 0.01, Spearman correlation coefficient = 0.78). The number of *Phaeomoniella* reads by HTAS was significantly higher in SWD (268.3 ± 54.8) than in MDW (94.8 ± 19.3) and AWD (58.6 ± 11.9) at t1, while at t2, significant differences were observed between SWD (199.6 ± 40.7) and MWD (141.8 ± 28.9) (Fig 7a). The number of copies of *P. chlamydospora* by ddPCR was significantly higher in SWD (42.8 ± 18.4) than in MDW (7.2 ± 4.3) and AWD (11.1 ± 5.4) at t1, while at t2, no significant differences were found between water regimes (Fig 7b).

**Fig. 7.**
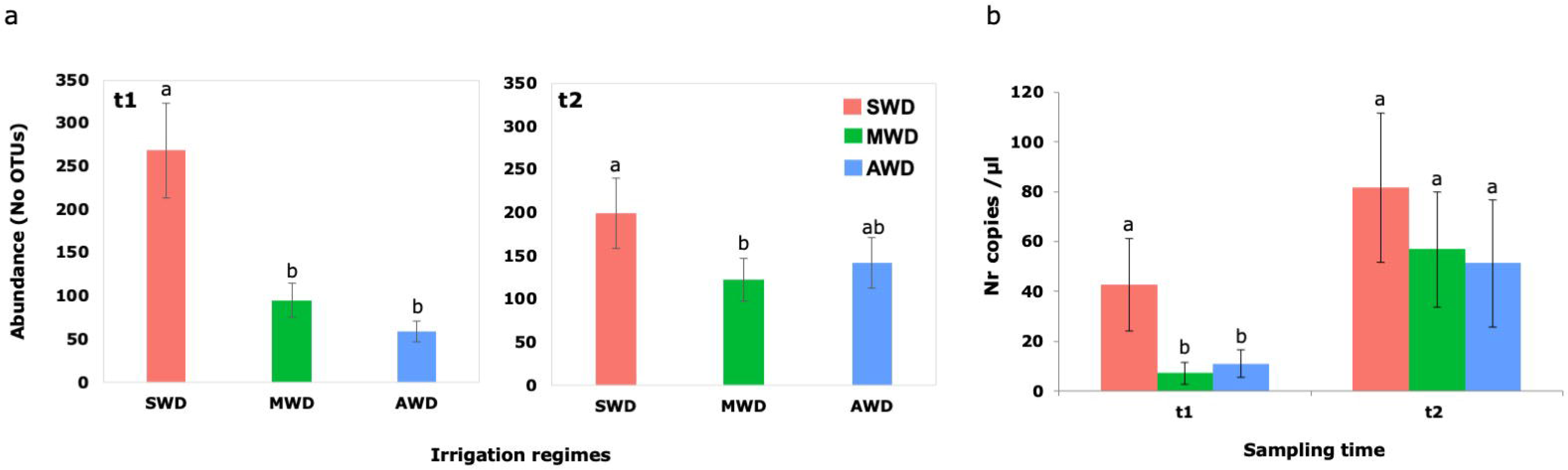
Abundance of OTUs (a) and DNA concentration (b) of *Phaeomoniella ch/amydospora* over two growing seasons (t1 and t2) across different irrigation regimes: Severe Water Deficit (SWD), Moderate Water Deficit (MWD), and No Water Deficit (AWD). Data represent mean of 24 replicates from High-Throughput Amplicon Sequencing and ddPCR analysis, with Standard Error of the Mean (SEM) error bars. Groups with different letters indicate significant differences per Tukey’s Honest Significant Difference Test.

### Water deficit reduces the complexity of co-occurrence networks among taxa

Before the experiment (t0), the xylem microbiome was complex, with a total of 386 interactions, of which 192 were positive and 194 negative (Fig. 8). At t2, we observed that plants with AWD were able to maintain a complex xylem microbiome with 325 interactions, of which 97 were positive and 228 were negative. On the contrary, plants with MWD had only 101 interactions, of which 94 were positive and 7 were negative. Plants under SWD presented only 64 interactions, of which 57 were positive and 7 were negative. (Fig. 8).

**Fig. 8.**
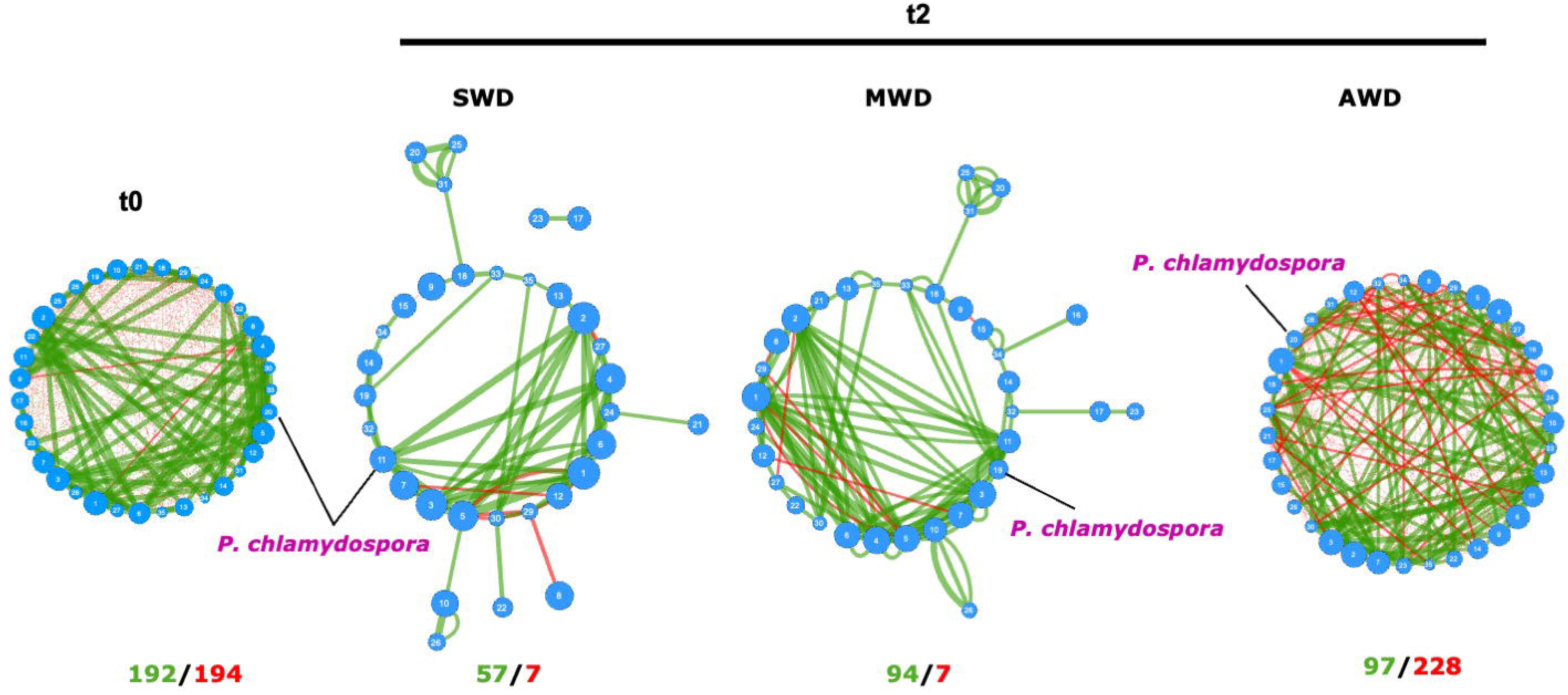
SparCC network map in operational taxonomic unit (OTU) level of the fungal xylem at baseline (tO) and after two growing seasons of applying three irrigation regimes: Severe Water Deficit (SWD), Moderate Water Deficit (MWD), and No Water Deficit (AWD). Each point represents a fungal OTU. Edges indicate the relationship between species. The size of the nodes follows betweenness centrality scores. The width of the edges follows SparCC correlation coefficients (green = positive correlation; red = negative correlation). The number of positive and negative correlations is shown below each network. Species with total relative abundance higher than 0.5% were chosen. Only statistically significant edges corresponding to correlations with a magnitude higher than 0.3 (*P* < 0.05) were drawn. The OTU corresponding to the genus Phaeomoniella chlamydospora is indicated in each network

## Discussion

Climate change poses a significant threat to water availability, particularly in Mediterranean regions, where grapevines are predominantly cultivated. The emergence of drought conditions not only impacts grapevine development but also exacerbates the susceptibility of stressed plants to various diseases by compromising their defense mechanisms. This study delves into the repercussions of induced water stress on the xylem-inhabiting fungal communities of grapevine, focusing on the primary pathogen associated with esca and Petri diseases, *P. chlamydospora*.

Our primary finding highlights that prolonged water stress not only altered the diversity but also the composition of the xylem fungal microbiome. Severe water deficit significantly reduced diversity compared to non-stressed plants. While water is essential for microbial growth, many microorganisms lack active water uptake mechanisms and depend on osmotically active substances in plant cells. Therefore, water deficit adversely affects the microbiomes of stressed plants. This aligns with prior research by Carbone et al. (2021), who reported reduced alpha-diversity in soil, rhizosphere, and root samples from grapevines subjected to medium and severe water stress.

Beyond diversity, water stress shaped the type of microbiome in grapevine xylem. A year of severe water stress notably increased the prevalence of pathotrophs in the xylem vessels, primarily comprising plant pathogens. After two years of water deficit, non-stressed plants exhibited significantly lower pathotroph abundance than their medium and severely stressed counterparts. Recent mathematical models predicting the impact of drought on disease development in the context of global warming corroborate our findings. Studies such as Songy et al. (2019) and Salinari et al. (2007), anticipated an escalation in pathogen presence and pathogenicity in environments experiencing water stress.

Our investigation, employing ddPCR and HTAS analyses, revealed that plants under severe water deficit exhibited a higher abundance of *P. chlamydospora* in the xylem compared to moderately stressed and non-stressed plants after two growing seasons. While the HTAS analysis focused on the genus level, given that *Phaeomoniella* comprises only one species, *P. chlamydospora* (Chen et al. 2022), a direct comparison with the ddPCR results was feasible. Bortolami et al. (2021) contradicted our results, demonstrating reduced abundance of *P. chlamydospora* and *Phaeocremonium minimum* in grapevines under drought stress. It is worth noting that their study collected samples from a single point on the perennial trunk of mature vines, focusing on the cultivar part. In contrast, our study obtained samples from three different points in rootstock cuttings of young plants. It is well established that the distribution of GTD pathogens is not uniform within plants (Aroca et al. 2010), and esca pathogens are often found in the rootstock section of mature plants (Elena et al. 2018). Therefore, collecting samples from only one point in the cultivar may underestimate the abundance of these fungi.

Several hypotheses can explain this increase in *P. chlamydospora* abundance in the xylem vessels of a young plant subjected to water stress. This abiotic factor in plants can weaken their defenses and compromise their ability to resist pathogen invasion (Gorshkov and Tsers, 2022). When a plant experiences water stress, it often prioritizes the allocation of limited resources towards essential physiological processes such as maintaining cell turgor pressure and photosynthesis, leaving fewer resources available for defense mechanisms against pathogens (Osakabe et al. 2014). Pathogenic fungi, like many other pathogens, take advantage of weakened plants as they may find it easier to penetrate the plant’s defenses and establish themselves within the xylem vessels (Doehlemann et al. 2017). Additionally, water stress can create favorable conditions for fungal growth by altering the plant’s physiology, such as increasing the concentration of sugars and other nutrients in the plant’s tissues, which can serve as a food source for the fungus (Bortolami et al. 2021). Water stress, as observed in our study, can also disrupt the balance of beneficial microorganisms in the plant’s xylem vessels, thereby potentially facilitating the colonization and proliferation of pathogenic fungi.

Another aspect worth noting from our research is that no leaf symptoms associated with young vine decline were observed in grafted plants during the two growing seasons. This concurs with earlier studies indicating that grapevine rootstock artificially inoculated with a high inoculum pressure of fungi associated with Petri disease may not exhibit visible symptoms one year after planting in the field (Gramaje et al. 2010). It is suggested that symptom expression may depend on the duration of the infection period (Graniti et al. 2000), necessitating longer assays to observe disease symptoms, and the amount of pathogen inoculum present in the xylem, referred to as the inoculum threshold (Hrycan et al. 2020). These vascular pathogens can usually act as latent pathogens and produce symptoms in the plant following other abiotic or biotic stress factors (Hrycan et al. 2020). In mature vines, Bortolami et al. (2021) observed a clear antagonism between drought and the appearance of leaf symptoms associated with esca. The authors postulated that drought might initiate systemic reactions that could disrupt pathogenic processes, such as diminishing fungal toxicity (Bortolami et al. 2021). Moreover, our study involved natural pathogen infections, resulting in a less concentrated inoculum compared to artificial inoculation, further complicating symptom expression and visualization.

According to the LEfSe analysis, our two growing seasons experiment revealed 15 genera significantly influencing the microbiome of the studied plants. Among these, *Trichoderma*, renowned for its biocontrol potential and already featured in commercial products against grapevine fungal diseases (Berbegal et al. 2020; Úrbez-Torres et al. 2020; Leal et al. 2021, 2023, 2024; Pollard-Flamand et al. 2022, 2023), and *Meyerozyma*, a potential biocontrol agent studied against various grapevine diseases (Alonso de Robador et al. 2023; Herrera-Balandrano et al. 2023), exhibit higher abundance in non-stressed plants compared to medium and severely stressed plants. Additionally, the genus *Mortierella* was more abundant in plants without water deficit. *Mortierella*, recognized as phosphate-solubilizing fungi, plays a crucial role in phosphorus cycling, contributing to increased plant weight and development (Osorio et al. 2001). Conversely, non-stressed plants showed a greater abundance of *Cadophora*, a pathogen responsible for Petri disease in grapevines (Gramaje et al. 2011; Úrbez-Torres et al. 2014; Maldonado-González et al. 2020).

Notably, severely stressed plants exhibited higher abundances of genera with biological control potential, such as *Acremonium*, known to enhance plant resistance against drought and temperature stress (White et al. 1992). Research by Llorens et al. (2019) demonstrated that the beneficial effect of *Acremonium* in wheat plants was associated with a reduced plant response to drought stress. While grapevines under water stress conditions seem to enlist a microbiome that enhances stress tolerance and provides protection against pathogens, SWD plants exhibited a notable alteration in their microbiome due to pathogens present in higher abundances compared to non-stressed plants. Examples of such pathogens include *Ceratobasidium* and *Phaeomoniella*.

Various beneficial and pathogenic microorganisms, including bacteria, fungi, and oomycetes, coexist in the rhizosphere alongside plants. In response to each other’s presence and the ever-changing environment, there is a complex web of communication among all parties involved (Venturi and Keel 2016; Ku et al. 2021). Consequently, our study delved into the co-occurrence networks within the xylem microbiome of plants under water deficit and non-stressed conditions, aiming to assess the impact of water deficit on fungal dynamics within the xylem microbiome. Water stress significantly altered the co-occurrence networks within the microbiome in the xylem. Plants subjected to a two-year water deficit exhibited fewer correlations between taxa, resulting in reduced network complexity, particularly under severe deficit conditions. The intricacy of the co-occurrence network plays a crucial role in maintaining network stability and enhancing resistance to external disturbances, including both biotic and abiotic stresses (Ding et al. 2023). This underscores the notion that water deficit can heighten grapevine susceptibility to a range of biotic and abiotic stresses.

## Conclusions

Our study underscored the profound impact of water availability, particularly in Mediterranean regions, in grapevine health. The induced water stress not only altered the diversity and composition of the fungal microbiome in the xylem vessels but also influenced co-occurrence networks, resulting in less complex networks, with less correlations between taxa, which increased the vulnerability of grapevines to various biotic and abiotic stresses. Severe water deficit significantly reduced microbial diversity, leading to a shift in the abundance of pathotrophs in the xylem, emphasizing the interconnectedness between water stress, microbiome dynamics, and plant health.

The investigation into *P. chlamydospora*, a key pathogen in Petri disease and esca, unraveled increased abundance under severe water deficit conditions. The combination of weakened plant defenses with altered physiological conditions, and changes in the surrounding microbial community might create an environment conducive to the increased abundance of *P. chlamydospora* in the xylem of young vines following water stress. The absence of visible symptoms in our study highlights the complexity of disease manifestation, urging longer assays and careful consideration of natural pathogen infections for comprehensive understanding.

In essence, our findings emphasize the intricate interplay between climate change, water stress, microbiome dynamics, and the health of grapevines. Addressing these complexities is crucial for developing effective strategies to mitigate the adverse effects on viticulture, ensuring the sustainability and resilience of grapevine cultivation in the face of a changing climate.

### Author contributions

MMM and DG conceived and designed the study. CL, MMM, MJC and DG wrote the manuscript. MMM and RB run the greenhouse experiment and carried out the sampling. TK, AE, MJC and DG performed the bioinformatics and statistical analyses. All authors read and approved the final manuscript.

### Funding

This study has been funded by the GLOBALVITI project under the Strategic Program for Consortia of National Business Research (CIEN) (Spanish Ministry of Economy, Industry and Competitiveness, Center for Industrial Technological Development (CDTI)).

### Availability of data and material

The datasets generated and analysed during the current study are available in the NCBI Sequence Read Archive (SRAs: SRR28044237 and SRR28046388) under the BioProject number PRJNA1078128. The OTU table, metadata and taxonomic classifications used in this study have been deposited in Figshare (ID: 197131).

The author(s) declare no conflict of interest.

## ACKNOWLEDGMENTS

We thank P. Yécora and M. Andrés-Sodupe for their assistance with sampling.

## Notes

### Competing Interest Statement

The authors have declared no competing interest.

## LITERATURE CITED

Abarenkov, K., Nilsson, R. H., Larsson, K. H., Alexander, I. J., Eberhardt, U., Erland, S., Hoiland, K., Kjoller, R., Larsson, E., Pennanen, T., Sen, R., Taylor, A., Tedersoo, L., Ursing, B., Vralstad, T., Liimatainen, K., Peintner, U., and Koljalg, U. 2010. The UNITE database for molecular identification of fungi–recent updates and future perspectives. New Phytol. 186:281–285.

Alonso de Robador, J. M., Ortega Pérez, N., Sanchez-Ballesta, M. T., Tello Mariscal, M. L., Pintos López, B., and Gómez-Garay, A. 2023. Plant defence induction by Meyerozyma guilliermondii in Vitis vinifera L. Agronomy. 13:2780.

Aroca, A., Gramaje, D., García-Jiménez, J., Armengol, J., and Raposo, R. 2010. Evaluation of the grapevine nursery process as a source of Phaeoacremonium spp. and Phaeomoniella chlamydospora and occurrence of trunk disease pathogens in rootstock mother vines in Spain. Eur. J. Plant Pathol. 126:165–174.

Aronesty, E. 2011. Ea-Utils: Command-Line Tools for Processing Biological Sequencing Data. [accessed 2022 Oct 15]. http://code.google.com/p/ea-utils.

Arora, N. K. 2019. Impact of climate change on agriculture production and its sustainable solutions. Environ. Sustain. 2(2):95–96.

Bengtsson-Palme, J., Ryberg, M., Hartmann, M., Branco, S., Wang, Z., Godhe, A., Wit, P., Sánchez-García, M., Ebersberger, I., Sousa, F., Amed, A., Jumpponen, A., Unterseher, M., Kristiansson, E., Abarenkov, K., Bertrand, Y., Sanli, K., Eriksson, M., Vik, U., Veldre, V., and Nilsson, H. 2013. Improved software detection and extraction of ITS1 and ITS2 from ribosomal ITS sequences of fungi and other eukaryotes for use in environmental sequencing. Methods Ecol. Evol. 4:914–919.

Berbegal, M., Ramón-Albalat, A., León, M., and Armengol, J. 2020. Evaluation of long-term protection from nursery to vineyard provided by Trichoderma atroviride SC1 against fungal grapevine trunk pathogens. Pest Manag. Sci. 76:967–977.

Bortolami, G., Gambetta, G. A., Delzon, S., Lamarque, L. J., Pouzoulet, J., Badel, E., Burlett, R., Charrier, G., Chocard, H., Dayer, S., Jansen, S., King, A., Lecomte, P., Lens, F., Torres-Ruiz, J. M., and Delmas, C. E. 2019. Exploring the hydraulic failure hypothesis of esca leaf symptom formation. Plant Physiol. 181(3):1163–1174.

Bortolami, G., Gambetta, G. A., Cassan, C., Dayer, S., Farolfi, E., Ferrer, N., Gibone, Y., Jolivet, J., Lecomte, P., and Delmas, C. E. 2021. Grapevines under drought do not express esca leaf symptoms. Proc. Natl. Acad. Sci. USA. 118(43):e2112825118.

Bortolami, G., Ferrer, N., Baumgartner, K., Delzon, S., Gramaje, D., Lamarque, L. J., Romanazzi, G., Gambetta, G. A., and Delmas, C. E. 2023. Esca grapevine disease involves lead hydraulic failure and represents a unique premature senescence process. Tree Physiol. 43:441–451.

Carbone, M., Alaniz, J., Mondino, P., Gelabert, M., Eichmeier, A., Tekielska, D., and Gramaje, D. 2021. Drought influences fungal community dynamics in the grapevine rhizosphere and root microbiome. J. Fungi. 7(9):686.

Chaves, M. M., Zarrouk, O., Francisco, R., Costa, J. M., Santos, T., Regalado, A. P., and Lopes, C. M. 2010. Grapevine under deficit irrigation: hints from physiological and molecular data. Ann. Bot. 105(5):661–676.

Chen, Q., Bakhshi, M., Balci, Y., Broders, K. D., Cheewangkoon, R., Chen, S. F., Fan, X. L., Gramaje, D., Halleen, F., Horta Jung, M., Jiang, N., Jung, T., Májek, T., Marincowitz, S., Milenković, I., Mostert, L., Nakashima, C., Nurul Faziha, I., Pan, M., Raza, M., Scanu, B., Spies, C. F. J., Suhaizan, L., Suzuki, H., Tian, C. M., Tomšovský, M., Úrbez-Torres, J. R., Wang, W., Wingfield, B. D., Wingfield, M. J., Yang, Q., Yang, X., Zare, R., Zhao, P., Groenewald, J. Z., Cai, L., and Crous, P. W. 2022. Genera of Phytopathogenic Fungi: GOPHY 4. Stud. Mycol. 101:417–564.

Ding, L., Tian, L., Li, J., Zhang, Y., Wang, M., and Wang, P. 2023. Grazing lowers soil multifunctionality but boosts soil microbial network complexity and stability in a subtropical grassland of China. Front. Microbiol. 13:1027097.

Doehlemann, G., Ökmen, B., Zhu, W., and Sharon, A. 2017. Plant Pathogenic Fungi. Microbiol. Spectr. 5(1).

Edgar, R. C. 2013. UPARSE: Highly accurate OTU sequences from microbial amplicon reads. Nat. Methods. 10:996–998.

Elena, G., Bruez, E., Rey, P., and Luque, J. 2018. Microbiota of grapevine Woody tissues with or without esca-foliar symptoms in northeast Spain. Phytopathol. Mediterr. 57:425–438.

Feng, K., Peng, X., Zhang, Z., Gu, S., He, Q., Shen, W., Wang, Z., Wang, D., Hu, Q., Li, Y., Wang, S., and Deng, Y. 2022. iNAP: an integrated network analysis pipeline for microbiome studies. iMeta. 1(2):e13.

Fischer, M., and Kassemeyer, H. H. 2012. Water regime and its possible impact on expression of Esca symptoms in Vitis vinifera: growth characters and symptoms in the greenhouse after artificial infection with Phaeomoniella chlamydospora. Vitis. 51(3):129–135.

Friedman, J., and Alm, E. J. 2012. Inferring correlation networks from genomic survey data. PLoS Comput. Biol. 8:e1002687.

Gambetta, G. A., Herrera, J. C., Dayer, S., Feng, Q., Hochberg, U., and Castellarin, S.D. 2020. The physiology of drought stress in grapevine: towards an integrative definition of drought tolerance. J. Exp. Bot. 71(16):4658–4676.

Gorshkov, V., and Tsers, I. 2022. Plant susceptible responses: the underestimated side of plant–pathogen interactions. Biol. Rev. 97:45–66.

Gramaje, D., and Armengol, J. 2011. Fungal trunk pathogens in the grapevine propagation process: potential inoculum sources, detection, identification, and management strategies. Plant Dis. 95:1040–1055.

Gramaje, D., Mostert, L., and Armengol, J. 2011. Characterization of Cadophora luteo- olivacea and C. melinii isolates obtained from grapevines and environmental samples from grapevine nurseries in Spain. Phytopathol. Mediterr. 50:S112–S126.

Gramaje, D., García-Jiménez, J., and Armengol, J. 2010. Field evaluation of grapevine rootstocks inoculated with fungi associated with Petri disease and esca. Am. J. Enol. Vitic. 61(4):512–520.

Gramaje, D., Eichmeier, A., Spetik, M., Carbone, M. J., Bujanda, R., Vallance, J., and Rey, P. 2022. Exploring the temporal dynamics of the fungal microbiome in rootstocks, the lesser-known half of the grapevine crop. J. Fungi. 8(5):421.

Gramaje, D., Úrbez-Torres, J. R., and Sosnowski, M. R. 2018. Managing grapevine trunk diseases with respect to etiology and epidemiology: current strategies and future prospects. Plant Dis. 102(1):12–39.

Graniti, A., Mugnai, L., and Surico, G. 2000. Esca of grapevine: A disease complex or a complex of diseases. Esca of Grapevine. 1000–1005.

Hacquard, S., Spaepen, S., Garrido-Oter, R., and Schulze-Lefert, P. 2017. Interplay between innate immunity and the plant microbiota. Annu. Rev. Phytopathol. 55:565–589.

Hardoim, P. R., Van Overbeek, L. S., Berg, G., Pirttilä, A. M., Compant, S., Campisano, A., and Sessitsch, A. 2015. The hidden world within plants: ecological and evolutionary considerations for defining functioning of microbial endophytes. Microbiol Mol. Biol. Rev. 79(3):293–320.

Herrera-Balandrano, D. D., Wang, S. Y., Wang, C. X., Shi, X. C., Liu, F. Q., and Laborda, P. 2023. Antagonistic mechanisms of yeasts Meyerozyma guilliermondii and M. caribbica for the control of plant pathogens: A review. Biol. Control. 105333.

Hrycan, J., Theilmann, J., Mahovlic, A., Boulé, J., and Úrbez-Torres, J. 2023. Health Status of Ready-to-Plant Grapevine Nursery Material in Canada Regarding Young Vine Decline Fungi. Plant Dis. 107:3708–3717.

Hrycan, J., Hart, M., Bowen, P., Forge, T., and Úrbez-Torres, J. R. 2020. Grapevine trunk disease fungi: their roles as latent pathogens and stress factors that favour disease development and symptom expression. Phytopathol. Mediterr. 59(3):395–424.

Jones, C. A., Jacobsen, J. S., and Wraith, J. M. 2005. Response of malt barley to phosphorus fertilization under drought conditions. J. Plant Nutr. 28(9):1605–1617.

Katoh, K., Asimenos, G., and Toh, H. 2009. Multiple Alignment of DNA Sequences with MAFFT. In: Posada D, editor. Bioinformatics for DNA Sequence Analysis. Methods Mol. Biol. 537:39–64. Totowa (NJ): Humana Press.

Ku, Y. S., Wang, Z., Duan, S., and Lam, H. M. 2021. Rhizospheric communication through mobile genetic element transfers for the regulation of microbe–plant interactions. Biology. 10(6):477.

Lamarque, L. J., Delmas, C. E., Charrier, G., Burlett, R., Dell’Acqua, N., Pouzoulet, J., and Delzon, S. 2023. Quantifying the grapevine xylem embolism resistance spectrum to identify varieties and regions at risk in a future dry climate. Sci. Rep. 13(1):7724.

Leal, C., Richet, N., Guise, J. F., Gramaje, D., Armengol, J., Fontaine, F., and Trotel- Aziz, P. 2021. Cultivar contributes to the beneficial effects of Bacillus subtilis PTA-271 and Trichoderma atroviride SC1 to protect grapevine against Neofusicoccum parvum. Front. Microbiol. 12:726132.

Leal, C., Gramaje, D., Fontaine, F., Richet, N., Trotel-Aziz, P., and Armengol, J. 2023. Evaluation of Bacillus subtilis PTA-271 and Trichoderma atroviride SC1 to control Botryosphaeria dieback and black-foot pathogens in grapevine propagation material. Pest Manag. Sci. 79(5):1674–1683.

Leal, C., Bujanda, R., López-Manzanares, B., Ojeda, S., Berbegal, M., Oneka, O., Gonzaga Santesteban, L., Palacios, J., and Gramaje, D. 2024. Evaluating treatments for the protection of grapevine pruning wounds from natural infection by trunk disease fungi. [Details Pending].

Lecomte, P., Darrieutort, G., Liminana, J. M., Comont, G., Muruamendiaraz, A., Legorburu, F. J., Choueiri, E., Jreijiri, F., El Amil, R., and Fermaud, M. 2012. New insights into esca of grapevine: the development of foliar symptoms and their association with xylem discoloration. Plant Dis. 96(7):924–934.

Liu, J., Zhao, F. L., Guo, Y., Fan, X. C., Wang, Y. J., and Wen, Y. Q. 2019. The ABA receptor-like gene VyPYL9 from drought-resistance wild grapevine confers drought tolerance and ABA hypersensitivity in Arabidopsis. Plant Cell Tissue Organ Cult. 138:543–558.

Llorens, E., Scalschi, L., Sharon, O., Vicedo, B., Sharon, A., and García-Agustín, P. 2022. Jasmonic acid pathway is required in the resistance induced by Acremonium sclerotigenum in tomato against Pseudomonas syringae. Plant Sci. 318:111210.

Lu, Y., Zhou, G., Ewald, J., Pang, Z., Shiri, T., and Xia, J. 2023. MicrobiomeAnalyst 2.0: comprehensive statistical, functional and integrative analysis of microbiome data. Nucleic Acids Res. 51:W310–W318.

Maldonado-González MM, Martinez-Diz MP, Andrés-Sodupe M, Bujanda R, Díaz- Losada E, Gramaje D. 2020. Quantification of Cadophora luteo-olivacea from grapevine nursery stock and vineyard soil using droplet digital PCR. Plant Dis. 104(8):2269–2274.

Martínez-Diz, M. P., Eichmeier, A., Spetik, M., Bujanda, R., Díaz-Fernández, Á., Díaz- Losada, E., and Gramaje, D. 2020. Grapevine pruning time affects natural wound colonization by wood-invading fungi. Fungal Ecol. 48:100994.

McLaren, M. R., and Callahan, B. J. 2020. Pathogen resistance may be the principal evolutionary advantage provided by the microbiome. Philos. Trans. R. Soc. Lond. B. Biol. Sci. 375(1808):20190592.

Mosedale, J. R., Abernethy, K. E., Smart, R. E., Wilson, R. J., and Maclean, I. M. 2016. Climate change impacts and adaptive strategies: lessons from the grapevine. Glob. Change Biol. 22(11):3814–3828.

Ngumbi, E., and Kloepper, J. 2016. Bacterial-mediated drought tolerance: current and future prospects. Appl. Soil Ecol. 105:109–125.

Nguyen, N. H., Song, Z., Bates, S. T., Branco, S., Tedersoo, L., Menke, J., and Kennedy, P. G. 2016. FUNGuild: an open annotation tool for parsing fungal community datasets by ecological guild. Fungal Ecol. 20:241–248.

O’Geen, A. T. 2013. Soil Water Dynamics. Nat. Educ. Knowl. 4(5):9.

Oliveira, M., Teles, J., Barbosa, P., Olazabal, F., and Queiroz, J. 2014. Shading of the fruit zone to reduce grape yield and quality losses caused by sunburn. OENO One. 48:179–187.

Osakabe, Y., Osakabe, K., Shinozaki, K., and Tran, L. P. 2014. Response of plants to water stress. Front. Plant Sci. 5:86.

Osorio, N. W., and Habte, M. 2001. Synergistic influence of an arbuscular mycorrhizal fungus and a P solubilizing fungus on growth and P uptake of Leucaena leucocephala in an Oxisol. Arid Land Res. Manag. 15(3):263–274.

Pérez-Harguindeguy, N., Diaz, S., Garnier, E., Lavorel, S., Poorter, H., Jaureguiberry, P., and Cornelissen, J. H. C. 2016. Corrigendum to: New handbook for standardised measurement of plant functional traits worldwide. Aust. J. Bot. 64(8):715–716.

Pineda, A., Kaplan, I., and Bezemer, T. M. 2017. Steering soil microbiomes to suppress aboveground insect pests. Trends Plant Sci. 22(9):770–778.

Pollard-Flamand, J., Boulé, J., Hart, M., and Úrbez-Torres, J. R. 2022. Biocontrol activity of Trichoderma species isolated from grapevines in British Columbia against Botryosphaeria dieback fungal pathogens. J. Fungi. 8:409.

Pollard-Flamand, J., Boulé, J., Hart, M., and Úrbez-Torres, J. R. 2023. Biological control of Botryosphaeria dieback of grapevine in British Columbia, Canada. Am. J. Enol. Vitic. 74:0740034.

R Core Team. 2023. R: A Language and Environment for Statistical Computing. Vienna (Austria): R Foundation for Statistical Computing. Available from: https://www.R-project.org/.

Reinhold-Hurek, B., Bünger, W., Sofía Burbano, C., Sabale, M., and Hurek, T. 2015. Roots shaping their microbiome: global hotspots for microbial activity. Annu. Rev. Phytopathol. 53:403–424.

Salinari, F., Giosue, S., Rossi, V., Tubiello, F. N., Rosenzweig, C., and Gullino, M. L. 2007. Downy mildew outbreaks on grapevine under climate change: elaboration and application of an empirical-statistical model. EPPO Bull. 37(2):317–326.

Scala, E., Micheli, M., Ferretti, F., Maresi, G., Zottele, F., and Scattolin, L. 2019. New diseases due to indigenous fungi in a changing world: The case of hop hornbeam canker in the Italian Alps. For. Ecol. Manag. 439:159–170.

Scholander, P. F., Bradstreet, E. D., Hemmingsen, E. A., and Hammel, H. T. 1965. Sap pressure in vascular plants: Negative hydrostatic pressure can be measured in plants. Science. 148:339–346.

Schultz, H. R., and Stoll, M. 2010. Some critical issues in environmental physiology of grapevines: future challenges and current limitations. Aust. J. Grape Wine Res. 16:4–24.

Segata, N., Izard, J., Waldron, L., Gevers, D., Miropolsky, L., Garrett, W. S., and Huttenhower, C. 2011. Metagenomic biomarker discovery and explanation. Genome Biol. 12:R60.

Shannon, P., Markiel, A., Ozier, O., Baliga, N. S., Wang, J. T., Ramage, D., and Ideker, T. 2003. Cytoscape: a software environment for integrated models of biomolecular interaction networks. Genome Res. 13(11):2498–2504.

Songy, A., Fernandez, O., Clément, C., Larignon, P., and Fontaine, F. 2019. Grapevine trunk diseases under thermal and water stresses. Planta. 249:1655–1679.

Sosnowski, M., Ayres, M., and Scott, E. 2016. Trunk diseases: the influence of water deficit on grapevine trunk disease. Wine Vitic. J. 31(4).

Turenne, C. Y., Sanche, S. E., Hoban, D. J., Karlowsky, J. A., and Kabani, A. M. 1999. Rapid identification of fungi by using the ITS2 genetic region and an automated fluorescent capillary electrophoresis system. J. Clin. Microbiol. 37:1846–1851.

Úrbez-Torres, J. R., Haag, P., Bowen, P., and O’Gorman, D. T. 2014. Grapevine trunk diseases in British Columbia: Incidence and characterization of the fungal pathogens associated with esca and Petri diseases of grapevine. Plant Dis. 98:469–482.

Úrbez-Torres, J. R., Tomaselli, E., Pollard-Flamand, J., Boule, J., Gerin, D., and Pollastro, S. 2020. Characterization of Trichoderma isolates from southern Italy, and their potential biocontrol activity against grapevine trunk disease fungi. Phytopathol. Mediterr. 59(3):425–439.

Van Leeuwen, C., and Darriet, P. 2016. The impact of climate change on viticulture and wine quality. J. Wine Econ. 11(1):150–167.

Venturi, V., and Keel, C. 2016. Signaling in the rhizosphere. Trends Plant Sci. 21:187–198.

Vetrovsky, T., Baldrian, P., and Morais, D. 2018. SEED 2: A user-friendly platform for amplicon high-throughput sequencing data analyses. Bioinformatics. 34:2292–2294.

White, R. H., Engelke, M. C., Morton, S. J., Johnson-Cicalese, J. M., and Ruemmele, B. A. 1992. Acremonium endophyte effects on tall fescue drought tolerance. Crop Sci. 32(6):1392–1396.

White, T. J., Bruns, T., Lee, S. H., and Taylor, J. W. 1990. Amplification and direct sequencing of fungal ribosomal RNA genes for phylogenetics. In: PCR Protocols: A Guide to Methods and Applications. San Diego (CA): Academic Press. p. 315–322.

