## Supplementary material for "Drought influences the fungal community structure, diversity, and functionality inhabiting the grapevine xylem and enhances the abundance of *Phaeomoniella chlamydospora*"

**Table S1.** Number of reads, total OTUs and alpha diversity indices.

| Index | | | | | | |
|---|---|---|---|---|---|---|
| | t1 | | | t2 | | |
| | SWD[a] | MWD[b] | AWD[c] | SWD | MWD | AWD |
| Reads | 10785.3 ± 117.4[d] | 12208.25 ± 478.2 | 15105.3 ± 1976.1 | 10710.1 ± 90.5 | 11408.2 ± 282.3 | 12809.0 ± 989.3 |
| OTUs | 107 | 106 | 115 | 96 | 104 | 122 |
| Chao1 | 21.61 ± 1.07 | 21.92 ± 1.98 | 24.11 ± 1.07 | 17.02 ± 1.41 | 18.57 ± 0.48 | 20.02 ± 0.45 |
| Shannon | 2.22 ± 0.03 | 2.19 ± 0.10 | 2.31 ± 0.08 | 1.91 ± 0.06 | 1.99 ± 0.04 | 1.97 ± 0.03 |

[a] Severe Water Deficit

[b] Moderate Water Deficit

[c] No Water Deficit

[d] Values are the mean of 24 replicates

**Table S2.** SparCC correlation analysis at genus level in the xylem vessels of plants after one (t1) and two (t2) growing seasons

t1

| Taxon1 | Taxon2 | Correlation | P.Value |
|---|---|---|---|
| Arthrobotrys | Aspergillus | 0.3076 | 0.0099 |
| Arthrobotrys | Phaeoacremonium | 0.3013 | 0.0198 |
| Aspergillus | Arthrobotrys | 0.3076 | 0.0099 |
| Butlerelfia | Monocillium | 0.3353 | 0.0198 |
| Cadophora | Monocillium | 0.5008 | 0.0099 |
| Cadophora | Penicillium | -0.3072 | 0.0495 |
| Cadophora | Phaeomoniella | -0.3758 | 0.0297 |
| Cadophora | Sakaguchia | -0.3104 | 0.0099 |
| Castanediella | Coniolariella | -0.3269 | 0.0099 |
| Coniolariella | Castanediella | -0.3269 | 0.0099 |
| Coniolariella | Galerina | 319 | 0.0198 |
| Coniolariella | Mortierella | -0.3779 | 0.0099 |
| Cyphellophora | Rhodotorula | 318 | 0.0198 |
| Dactylonectria | Monocillium | 0.3182 | 0.0297 |
| Exophiala | Humicola | 0.3268 | 0.0198 |
| Filobasidium | Sugiyamaella | 0.3438 | 0.0099 |
| Galerina | Coniolariella | 319 | 0.0198 |
| Humicola | Exophiala | 0.3268 | 0.0198 |
| Lyomyces | Phaeoacremonium | 0.3017 | 0.0099 |
| Lyomyces | Sugiyamaella | 0.3018 | 0.0099 |
| Malassezia | Minimelanolocus | 0.3668 | 0.0099 |
| Metarhizium | Solicoccozyma | 0.41 | 0.0099 |
| Metarhizium | Wallemia | 0.3305 | 0.0099 |
| Meyerozyma | Penicillium | -0.3071 | 0.0396 |
| Meyerozyma | Psathyrella | 0.3192 | 0.0099 |
| Minimelanolocus | Malassezia | 0.3668 | 0.0099 |
| Monocillium | Butlerelfia | 0.3353 | 0.0198 |
| Monocillium | Cadophora | 0.5008 | 0.0099 |
| Monocillium | Dactylonectria | 0.3182 | 0.0297 |
| Monocillium | Neophysalospora | -0.3735 | 0.0297 |
| Monocillium | Sakaguchia | -0.3628 | 0.0297 |
| Monocillium | Talaromyces | 325 | 0.0198 |
| Monocillium | unknown | 331 | 0.0396 |
| Mortierella | Coniolariella | -0.3779 | 0.0099 |
| Mortierella | Naganishia | -0.3902 | 0.0099 |
| Mortierella | Penicillium | 0.4649 | 0.0099 |
| Mortierella | Psathyrella | 0.3218 | 0.0099 |
| Mortierella | Solicoccozyma | 409 | 0.0099 |
| Naganishia | Mortierella | -0.3902 | 0.0099 |
| Naganishia | Wallemia | -0.3097 | 0.0198 |
| Neoanthostomella | Psathyrella | 0.3337 | 0.0099 |
| Neoanthostomella | Rosellinia | 0.3422 | 0.0099 |
| Neoanthostomella | unknown | -311 | 0.0396 |
| Neophysalospora | Monocillium | -0.3735 | 0.0297 |
| Penicillium | Cadophora | -0.3072 | 0.0495 |
| Penicillium | Meyerozyma | -0.3071 | 0.0396 |
| Penicillium | Mortierella | 0.4649 | 0.0099 |
| Penicillium | Solicoccozyma | 0.4332 | 0.0099 |
| Penicillium | Thelonectria | 0.3826 | 0.0099 |
| Phaeoacremonium | Arthrobotrys | 0.3013 | 0.0198 |
| Phaeoacremonium | Lyomyces | 0.3017 | 0.0099 |
| Phaeomoniella | Cadophora | -0.3758 | 0.0297 |
| Phaeomoniella | Phlogicylindrium | 0.3206 | 0.0099 |
| Phaeomoniella | Psathyrella | -0.3116 | 0.0099 |
| Phlogicylindrium | Phaeomoniella | 0.3206 | 0.0099 |
| Phlogicylindrium | Sporothrix | 0.4915 | 0.0099 |
| Psathyrella | Meyerozyma | 0.3192 | 0.0099 |
| Psathyrella | Mortierella | 0.3218 | 0.0099 |
| Psathyrella | Neoanthostomella | 0.3337 | 0.0099 |
| Psathyrella | Phaeomoniella | -0.3116 | 0.0099 |
| Psathyrella | Talaromyces | 0.4277 | 0.0099 |
| Rhodotorula | Cyphellophora | 318 | 0.0198 |
| Rosellinia | Neoanthostomella | 0.3422 | 0.0099 |
| Sakaguchia | Cadophora | -0.3104 | 0.0099 |
| Sakaguchia | Monocillium | -0.3628 | 0.0297 |
| Solicoccozyma | Metarhizium | 0.41 | 0.0099 |
| Solicoccozyma | Mortierella | 409 | 0.0099 |
| Solicoccozyma | Penicillium | 0.4332 | 0.0099 |
| Sporothrix | Phlogicylindrium | 0.4915 | 0.0099 |
| Sporothrix | Talaromyces | -0.3909 | 0.0099 |
| Sugiyamaella | Filobasidium | 0.3438 | 0.0099 |
| Sugiyamaella | Lyomyces | 0.3018 | 0.0099 |
| Talaromyces | Monocillium | 325 | 0.0198 |
| Talaromyces | Psathyrella | 0.4277 | 0.0099 |
| Talaromyces | Sporothrix | -0.3909 | 0.0099 |
| Thelonectria | Penicillium | 0.3826 | 0.0099 |
| Thelonectria | Wallemia | 0.3258 | 0.0198 |
| unknown | Monocillium | 331 | 0.0396 |
| unknown | Neoanthostomella | -311 | 0.0396 |
| Wallemia | Metarhizium | 0.3305 | 0.0099 |
| Wallemia | Naganishia | -0.3097 | 0.0198 |
| Wallemia | Thelonectria | 0.3258 | 0.0198 |

t2

| Taxon1 | Taxon2 | Correlation | P.Value |
|---|---|---|---|
| Aspergillus | Calyptosphaeria | -0.2583 | 0.0297 |
| Aspergillus | Malassezia | 0.3046 | 0.0198 |
| Cadophora | Solicoccozyma | 0.2354 | 0.0198 |
| Cadophora | Sporothrix | -0.2671 | 0.0297 |
| Calyptosphaeria | Aspergillus | -0.2583 | 0.0297 |
| Calyptosphaeria | Minimelanolocus | 0.2469 | 0.0297 |
| Candida | Cyphellophora | -0.2473 | 0.0396 |
| Candida | unknown | -0.2613 | 0.0396 |
| Candida | Xylaria | -0.2438 | 0.0495 |
| Ceratobasidium | Mortierella | -0.2565 | 0.0198 |
| Ceratobasidium | Phaeomoniella | 0.3077 | 0.0297 |
| Ceratobasidium | Psathyrella | -0.2908 | 0.0099 |
| Ceratobasidium | Remispora | -0.2313 | 0.0297 |
| Ceratobasidium | Rhodotorula | 0.2077 | 0.0495 |
| Ceratobasidium | unknown | -0.2756 | 0.0297 |
| Coniolariella | Meyerozyma | -0.2074 | 0.0495 |
| Cyphellophora | Candida | -0.2473 | 0.0396 |
| Cyphellophora | Psathyrella | -0.2451 | 0.0099 |
| Exophiala | Xylaria | -0.2895 | 0.0198 |
| Malassezia | Aspergillus | 0.3046 | 0.0198 |
| Malassezia | Monocillium | 0.5109 | 0.0198 |
| Malassezia | Mortierella | -0.2981 | 0.0099 |
| Malassezia | Penicillium | 0.6006 | 0.0099 |
| Malassezia | Wallemia | 0.5287 | 0.0495 |
| Meira | Monocillium | -0.3037 | 0.0396 |
| Meira | Mortierella | 0.2266 | 0.0198 |
| Meira | Penicillium | -0.3056 | 0.0297 |
| Meyerozyma | Coniolariella | -0.2074 | 0.0495 |
| Meyerozyma | Xylaria | 0.2657 | 0.0099 |
| Minimelanolocus | Calyptosphaeria | 0.2469 | 0.0297 |
| Minimelanolocus | Phlogicylindrium | 0.3355 | 0.0099 |
| Monocillium | Malassezia | 0.5109 | 0.0198 |
| Monocillium | Meira | -0.3037 | 0.0396 |
| Monocillium | Mortierella | -494 | 0.0099 |
| Monocillium | Penicillium | 637 | 0.0297 |
| Mortierella | Ceratobasidium | -0.2565 | 0.0198 |
| Mortierella | Malassezia | -0.2981 | 0.0099 |
| Mortierella | Meira | 0.2266 | 0.0198 |
| Mortierella | Monocillium | -494 | 0.0099 |
| Mortierella | Psathyrella | 0.2256 | 0.0297 |
| Mortierella | Rosellinia | -0.2371 | 0.0495 |
| Penicillium | Malassezia | 0.6006 | 0.0099 |
| Penicillium | Meira | -0.3056 | 0.0297 |
| Penicillium | Monocillium | 637 | 0.0297 |
| Penicillium | Xylaria | 0.2836 | 0.0396 |
| Phaeoacremonium | Solicoccozyma | -0.2062 | 0.0396 |
| Phaeoacremonium | Talaromyces | 0.2314 | 0.0495 |
| Phaeomoniella | Ceratobasidium | 0.3077 | 0.0297 |
| Phaeomoniella | Phlogicylindrium | -0.2954 | 0.0198 |
| Phaeomoniella | Trichoderma | -0.2115 | 0.0396 |
| Phaeomoniella | unknown | -0.2426 | 0.0297 |
| Phaeomoniella | Wallemia | 0.4279 | 0.0396 |
| Phlogicylindrium | Minimelanolocus | 0.3355 | 0.0099 |
| Phlogicylindrium | Phaeomoniella | -0.2954 | 0.0198 |
| Psathyrella | Ceratobasidium | -0.2908 | 0.0099 |
| Psathyrella | Cyphellophora | -0.2451 | 0.0099 |
| Psathyrella | Mortierella | 0.2256 | 0.0297 |
| Psathyrella | Remispora | 0.2114 | 0.0396 |
| Remispora | Ceratobasidium | -0.2313 | 0.0297 |
| Remispora | Psathyrella | 0.2114 | 0.0396 |
| Rhodotorula | Ceratobasidium | 0.2077 | 0.0495 |
| Rhodotorula | Solicoccozyma | 0.2486 | 0.0396 |
| Rosellinia | Mortierella | -0.2371 | 0.0495 |
| Sakaguchia | Trichoderma | -0.2188 | 0.0396 |
| Solicoccozyma | Cadophora | 0.2354 | 0.0198 |
| Solicoccozyma | Phaeoacremonium | -0.2062 | 0.0396 |
| Solicoccozyma | Rhodotorula | 0.2486 | 0.0396 |
| Solicoccozyma | Talaromyces | -0.2782 | 0.0099 |
| Sporothrix | Cadophora | -0.2671 | 0.0297 |
| Sporothrix | Talaromyces | 0.2032 | 0.0495 |
| Talaromyces | Phaeoacremonium | 0.2314 | 0.0495 |
| Talaromyces | Solicoccozyma | -0.2782 | 0.0099 |
| Talaromyces | Sporothrix | 0.2032 | 0.0495 |
| Trichoderma | Phaeomoniella | -0.2115 | 0.0396 |
| Trichoderma | Sakaguchia | -0.2188 | 0.0396 |
| unknown | Candida | -0.2613 | 0.0396 |
| unknown | Ceratobasidium | -0.2756 | 0.0297 |
| unknown | Phaeomoniella | -0.2426 | 0.0297 |
| Wallemia | Malassezia | 0.5287 | 0.0495 |
| Wallemia | Phaeomoniella | 0.4279 | 0.0396 |
| Xylaria | Candida | -0.2438 | 0.0495 |
| Xylaria | Exophiala | -0.2895 | 0.0198 |
| Xylaria | Meyerozyma | 0.2657 | 0.0099 |
| Xylaria | Penicillium | 0.2836 | 0.0396 |

**Table S3.** Relative proportion (%) of fungal function from grapevine xylem at each irrigation regime inferred by FunGuild.

| | Year 1 | | | Year 2 | | |
|---|---|---|---|---|---|---|
| | SWD[a] | MWD[b] | AWD[c] | SWD | MWD | AWD |
| Pathotroph | 43.5 ± 3.3 Aa | 28.1 ± 0.9 Ab | 28.4 ± 0.5 Ab | 27.8 ± 3.2 Aa | 29.1 ± 0.9 Aa | 15.5 ± 1.1 Bb |
| Saprotroph | 24.2 ± 5.1 Ba | 31.0 ± 1.3 Aa | 35.0 ± 2.1 Aa | 31.5 ± 1.9 Ab | 43.9 ± 2.8 Aa | 45.9 ± 2.1 Aa |
| Symbiotroph | 25.5 ± 2.2 Ba | 34.8 ± 0.5 Aa | 30.9 ± 2.1 Aa | 31.7 ± 2.5 Aa | 19.8 ± 3.2 Aa | 29.4 ± 2.1 ABa |
| Unassigned | 6.8 ± 0.4 Ba | 6.0 ± 0.4 Ba | 5.6 ± 1.6 Ba | 8.9 ± 0.5 Ba | 7.2 ± 1.4 Aa | 9.0 ± 0.9 Ba |

[a] Severe Water Deficit

[b] Moderate Water Deficit

[c] No Water Deficit

Tukey's test at $P < 0.05$ level. Means followed by the same letter do not differ significantly ($P < 0.05$). Capital letters are for comparison of means among functional groups within each irrigation regime. Small letters are for comparison of means among irrigation regimes within each functional group.

**Table S4.** Compositions and relative abundance (%) of fungal functional groups (guild) inferred by FunGuild .

| | | Year 1 | | | Year 2 | | |
|---|---|---|---|---|---|---|---|
| | | **SWD**[a] | **MDW**[b] | **AWD**[c] | **SWD** | **MWD** | **AWD** |
| Pathotroph | Plant Pathogen | 31.2 Aa | 18.3 ABab | 12.3 Bb | 15.6 BCab | 18.9 Aa | 8.6 CDb |
| | Animal Pathogen | 4.3 Ba | 3.1 Ca | 7.7 BCa | 4.5 Da | 5.3 Ba | 3.4 Da |
| | Fungal Parasite | 3.4 Ba | 3.2 Ca | 3.3 Ca | 2.2 Da | 0.9 Ba | 1.6 Da |
| | Lichen Parasite | 4.5 Ba | 3.5 Ca | 5.1 Ca | 4.8 Da | 3.8 Ba | 1.9 Da |
| | Undefined Parasite | 0.1 Ba | 0 Ca | 0 Ca | 0.7 Da | 0.2 Ba | 0 Da |
| Saprotroph | Soil Saprotroph | 4.3 Ba | 5.2 Ca | 4.5 Ca | 6.7 CDa | 8.7 Ba | 9.7 CDa |
| | Wood Saprotroph | 3.4 Ba | 5.6 Ca | 5.6 Ca | 5.6 Db | 12.4 ABab | 15.6 BCa |
| | Dung Saprotroph | 4.7 Ba | 1.5 Ca | 3.7 Ca | 3.3 Da | 4.6 Ba | 3.3 Da |
| | Plant Saprotroph | 3.5 Ba | 6.2 Ca | 5.8 Ca | 3.7 Da | 2.1 Ba | 3.0 Da |
| | Litter Saprotroph | 1.6 Ba | 1.2 Ca | 3.3 Ca | 0.9 Da | 1.2 Ba | 3.7 Da |
| | Undefined Saprotroph | 6.7 Ba | 11.3 BCa | 12.2 Ba | 11.3 BC | 14.9 AB | 10.7 CD |
| Symbiotroph | Endophyte | 16.7 ABa | 25.1 Aa | 25.6 Aa | 26.4 Aa | 14.1 ABa | 26.7 Aa |
| | Epiphyte | 0 Ba | 0.3 Ca | 0.5 Ca | 1.2 Da | 0.3 Ba | 0.3 Da |
| | Endomycorrhizal | 7.6 Ba | 9.4 BCa | 4.8 Ca | 3.9 Da | 5.3 Ba | 2.6 Da |
| | Ectomycorrhizal | 0 Ba | 0.1 Ca | 0 Ca | 0 Da | 0 Ba | 0 Da |
| | Arbuscular Mycorrhizal | 1.2 Ba | 0 Ca | 0 Ca | 0.1 Da | 0 Ba | 0 Da |
| Unassigned | | 6.8 Ba | 6.0 Ca | 5.6 Ca | 8.9 CDa | 7.2 Ba | 9.1 CDa |

[a] Severe Water Deficit

[b] Moderate Water Deficit

[c] No Water Deficit

Tukey's test at *P* < 0.05 level. Means followed by the same letter do not differ significantly (*P*< 0.05). Capital letters are for comparisons of means of functional groups within each irrigation regime and year. Small letters are for comparison of means of each functional group among irrigation regimes within each year.

**Fig. S1**
Measurements of Stomatal conductance (A) and Predawn Leaf Water Potential (B) during the first (t0 to t1) and second (t1 to t2) growing seasons.

**A**

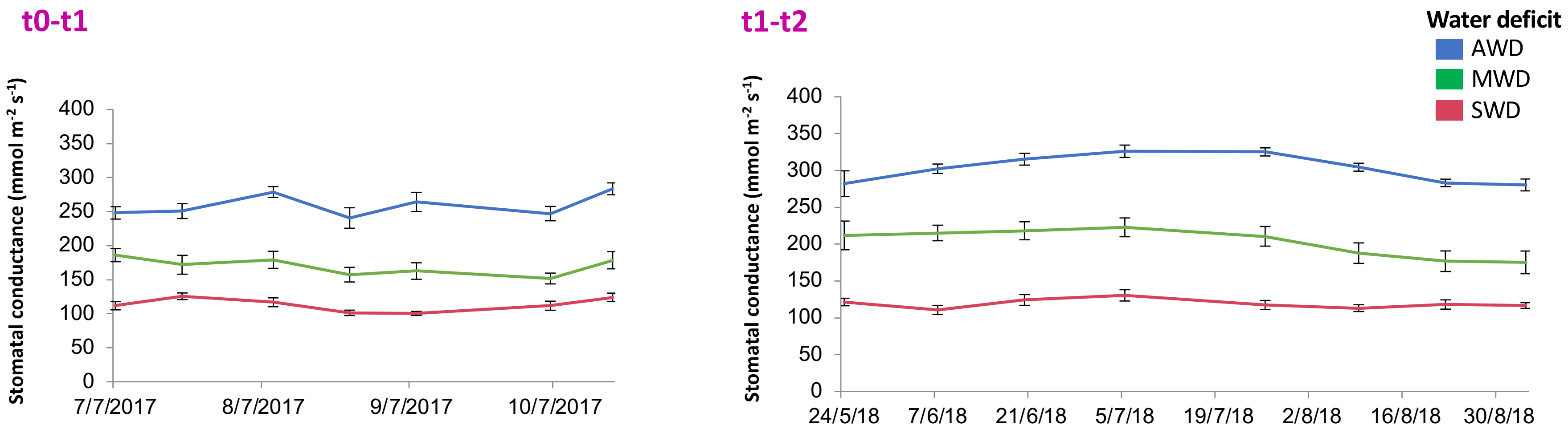


**B**

**t0-t1**

| Water potential (Mpa) | | | | | | |
|---|---|---|---|---|---|---|
| Water Deficit | Leaves | | | | | Mean |
| | 1 | 2 | 3 | 4 | 5 | |
| AWD | -0.7 | -0.6 | -0.6 | -0.6 | -0.7 | -0.64 |
| MWD | -0.9 | -0.8 | -0.8 | -0.9 | -0.8 | -0.84 |
| SWD | -0.9 | -0.75 | -0.75 | -0.8 | -0.85 | -0.81 |

**t1-t2**

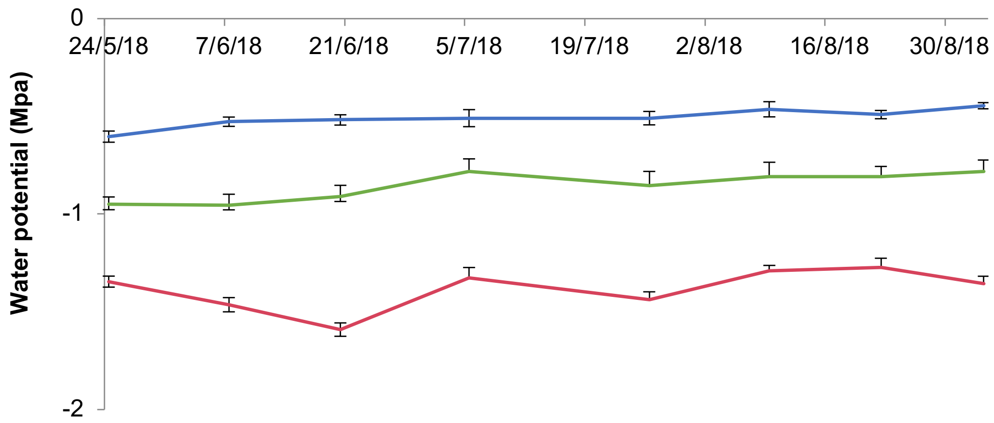

**Fig. S2**
Shoot weight of 'Tempranillo' grapevines grafted onto '110 Richter' rootstock under different irrigation regimes: 25% field capacity (severe water deficit, SWD), 50% field capacity (moderate water deficit, MWD), and 100% field capacity (no water deficit, AWD). Data represent mean values of two sets of 12 grafted plants each. Error bars denote standard error of the mean (SEM). Groups not sharing the same letter differ significantly according to Tukey's Honest Significant Difference Test.

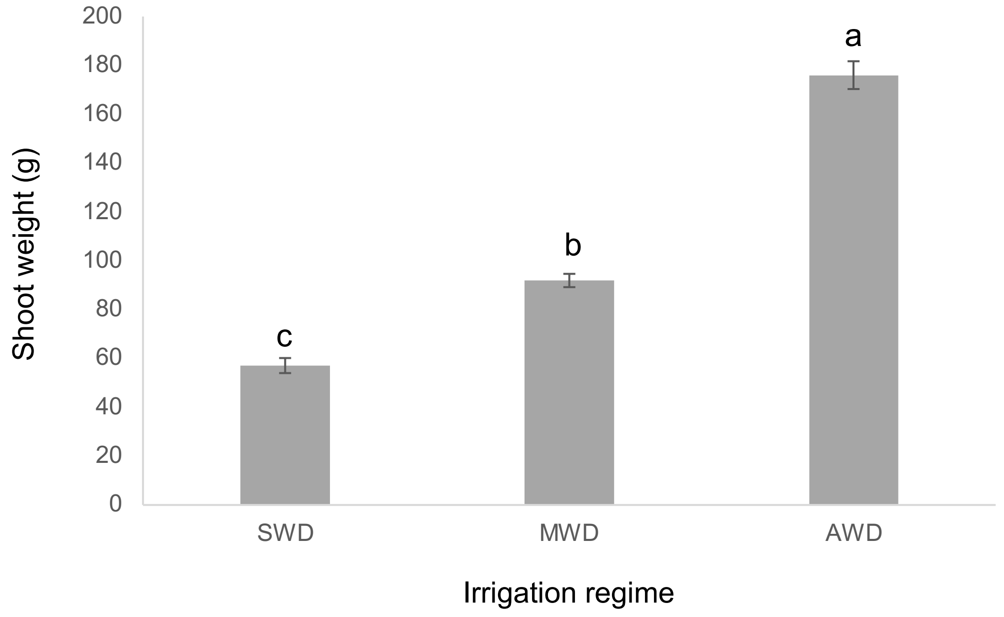

**Fig. S3**
Rarefaction curve values for each sample.

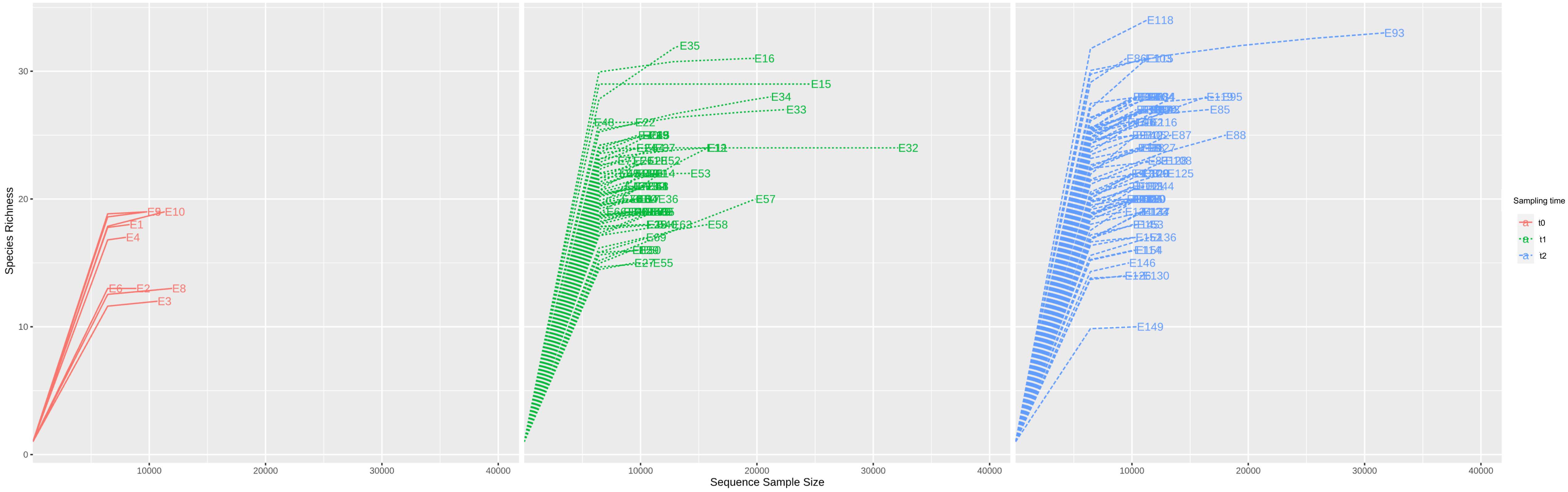

**Fig. S4**
Alpha diversity of fungi in the xylem microbiome following one (t1) and two (t2) growing seasons. *P*-values for each diversity index are indicated within the graphs.

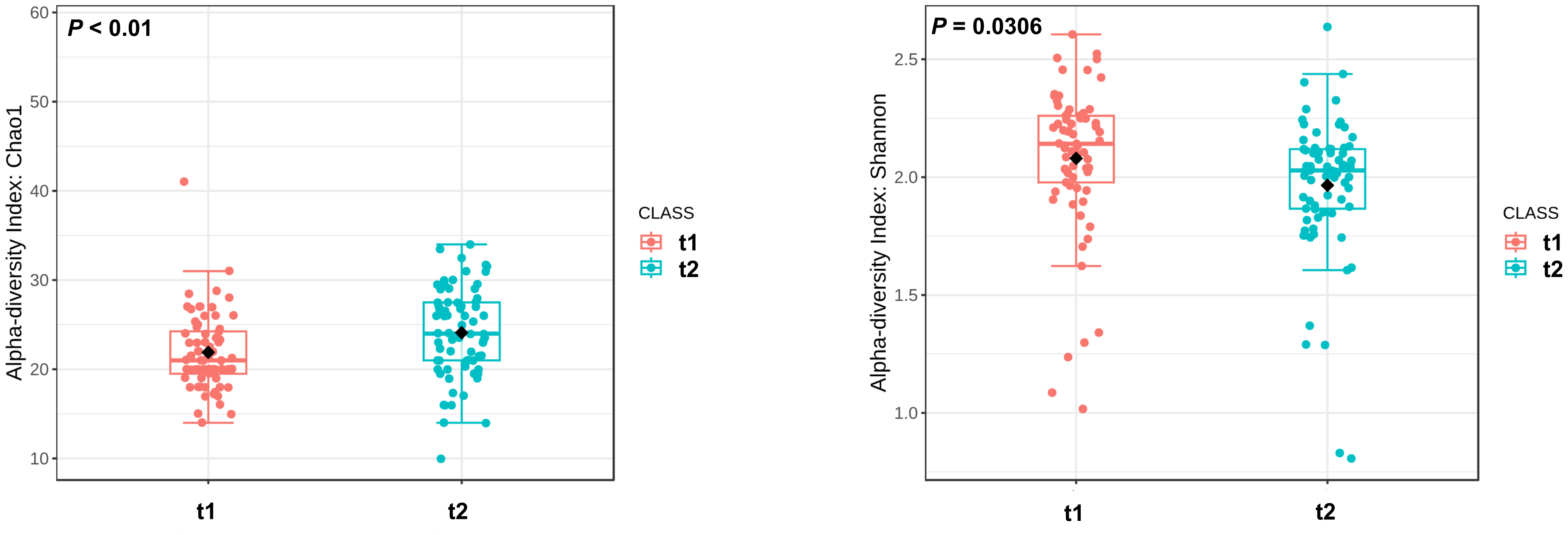
